# Inferring biophysical models of evolution from genome-wide patterns of codon usage

**DOI:** 10.1101/578815

**Authors:** Willow B. Kion-Crosby, Michael Manhart, Alexandre V. Morozov

## Abstract

Frequencies of synonymous codons are typically non-uniform, despite the fact that such codons correspond to the same amino acid in the genetic code. This phenomenon, known as codon bias, is broadly believed to be due to a combination of factors including genetic drift, mutational biases, and selection for speed and accuracy of codon translation; however, quantitative modeling of codon bias has been elusive. We have developed a biophysical population genetics model which explains genome-wide codon frequencies observed across 20 organisms. We assume that codons evolve independently of each other under the influence of mutation and selection forces, and that the codon population has reached evolutionary steady state. Our model implements codon-level treatment of mutations with transition/transversion biases, and includes two contributions to codon fitness which describe codon translation speed and accuracy. Furthermore, our model includes wobble pairing – the possibility of codon-anticodon base pairing mismatches at the 3’ nucleotide position of the codon. We find that the observed patterns of genome-wide codon usage are consistent with a strong selective penalty for mistranslated amino acids. Thus codons undergo purifying selection and their relative frequencies are affected in part by mutational robustness. We find that the dependence of codon fitness on translation speed is weaker on average compared to the strength of selection against mistranslation. Although no constraints on codon-anticodon pairing are imposed *a priori*, a reasonable hierarchy of pairing rates, which conforms to the wobble hypothesis and is consistent with available structural evidence, emerges spontaneously as a model prediction. Finally, treating the translation process explicitly in the context of a finite ribosomal pool has allowed us to estimate mutation rates per nucleotide directly from the coding sequences. Reminiscent of Drake’s observation that mutation rates are inversely correlated with the genome size, we predict that mutation rates are inversely proportional to the number of genes. Overall, our approach offers a unified biophysical and population genetics framework for studying codon bias across all domains of life.

## Introduction

The central dogma of molecular biology states that consecutive triplets of nucleotides called codons are translated into amino acids during protein production (Alberts *et al.*, 2015; Crick, 1958). As there are 64 codons and 20 amino acids, the translation code is degenerate, with as many as 6 codons translated into a single amino acid. Pronounced differences in synonymous codon usage are observed in any organism for which protein coding sequences are available and therefore codon frequencies can be reliably computed. These genome-wide differences are known as codon bias (Hershberg and Petrov, 2008; Napolitano *et al.*, 2016; Nielsen *et al.*, 2007; Sharp *et al.*, 2005, 2010). Since codon usage is one of the most fundamental features of genomes, a quantitative understanding of its evolution is critical to molecular biology.

Because a protein’s function is determined solely by its amino acid sequence, arguably the most basic mechanism for dictating the choice of synonymous codons is neutral evolution on a fitness landscape shaped by selective penalties for amino acid mistranslation (Akashi, 2001; Kimura, 1981). In this approach, non-uniform codon frequencies are produced due to mutational robustness (van Nimwegen *et al.*, 1999) and transition/transversion mutational biases (Yang, 2006).

Another popular explanation for the global codon bias involves selection and postulates that certain codons are translated more efficiently than others, resulting in higher protein production rates and therefore higher cellular growth rates or fitness (Akashi, 2001; Bulmer, 1991; Duret, 2002; Plotkin and Kudla, 2011). This translation efficiency can be characterized as a balance between translation speed and accuracy (Tuller *et al.*, 2010): a particular codon may be more rapidly translated due to a higher concentration of the corresponding tRNAs (a hypothesis supported by the correlation between tRNA gene copy numbers and codon frequencies (Kanaya *et al.*, 1999)), but may also cause more translation errors. The translation errors can be viewed through the lens of the wobble hypothesis, which states that each codon can be recognized by non-cognate tRNA species, with mispairings that occur at the 3’ nucleotide position in the codon (Crick, 1966; Stoletzki and Eyre-Walker, 2007).

Codon bias has been previously examined through population genetic models which incorporate mutation, selection, and drift in a system of two codon types (Bulmer, 1991; Kimura, 1991; Kondrashov, 1995; Li, 1987; McVean and Charlesworth, 1999). Since a complete treatment of a multi-allelic mutation-selection-drift model is prohibitively complex, especially in the polymorphic limit (Mustonen and Lässig, 2010), previous work has attributed the difference in codon frequencies to a balance between selection and drift, with mutations playing a subordinate role (Hershberg and Petrov, 2008). However, because selection strength has to be inversely proportional to the effective population size to reproduce the observed genomic codon frequencies, this approach leads to the “fine-tuning” problem in which selective advantages of the preferred codons have to vary through many orders of magnitude in order to reflect a broad range of effective population sizes (Charlesworth, 2009). It is challenging to provide a biophysical explanation for this behavior.

In contrast, our model focuses on the interplay between mutational and selective forces acting on individual codons: the observed codon frequencies emerge as a steady-state balance between mutational forces on one hand, and selection on translation speed and accuracy on the other. We have explicitly modeled the evolutionary process on the full 64-codon mutational network in a population of organisms whose fitness is determined by genomic codon content. Our approach is based on a realistic codon-level mutation model which includes transition/transversion biases and mutational robustness, and allows for non-cognate tRNA-mRNA pairings consistent with the wobble hypothesis. Using this selection-mutation framework, we were able to accurately predict genome-wide codon frequencies in a variety of organisms spanning both prokaryotic and eukaryotic domains. Our predictions of the codon-anticodon pairing rates are largely consistent with previously postulated wobble rules (Crick, 1966) and with the crystallographic analysis of wobble base pairs in the context of the ribosomal decoding center (Murphy IV and Ramakrishnan, 2004). We incorporated Bulmer’s biophysical model, which explicitly describes the details of the translation process given a finite ribosomal pool (Bulmer, 1991), into our approach, and estimated single-nucleotide mutation rates using biophysical model parameters such as ribosomal on-rates and codon translation times. Finally, our framework yields a potential explanation for Drake’s rule (Drake, 1991), which states that the mutation rate is inversely proportional to the genome size.

## Results

### Biophysical model of codon evolution

We have developed a biophysical model of codon population dynamics which is designed to predict genome-wide codon frequencies under the assumption that each organism in a population is subjected to mutation and selection on single-codon translation efficiency. We focus our attention primarily on the species with large effective codon population sizes, allowing us to neglect the effects of genetic drift in our model, which assumes that codons evolve independently at multiple genomic sites throughout the genome.

We consider the fitness of each organism, *w*, given the presence of a codon *c* at a particular genomic location and the optimal amino acid or STOP instruction *j* at that location, as the product of two terms modeling translation speed and accuracy, respectively (Materials and Methods):

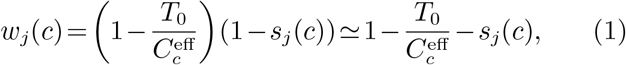

where 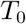 sets the overall scale of the selection coefficient in the first term, which penalizes for slow codon translation, and 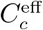 is the effective tRNA gene copy number. The approximation in Eq. (1) is valid when the two selection terms 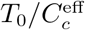 and *s*_*j*_(*c*) are small, as is generally expected for selection on a single codon. Since according to the wobble hypothesis non-cognate codon-anticodon pairing is allowed at the 3’ codon position, 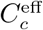 is computed as a weighted sum over all possible codon-anticodon pairings,

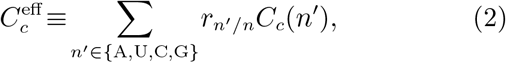

where *r*_*n′/n*_ is the codon-anticodon pairing rate associated with the nucleotide pairing *n′*/*n* at the 5’ anticodon and 3’ codon positions, respectively, and *C*_*c*_(*n′*) is the corresponding anticodon tRNA gene copy number, which we assume is proportional to the total number of tRNA molecules in the cell. For brevity, we shall refer to *r*_*n′/n*_ as “pairing rates” from now on. Note that the pairing rates are defined to be dimensionless and unnormalized (see Materials and Methods for details).

In the second term on the right-hand side of Eq. (1), *s*_*j*_(*c*) is the amino-acid-level selection coefficient which penalizes for incorrect amino acid translations due to wobble pairing:

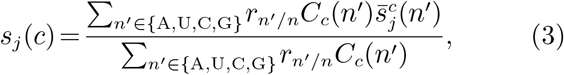

where 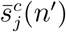 is either zero when the tRNA bound to codon *c* is charged with the optimal amino acid *j*, or a constant penalty, *s*, for any other amino acid. Thus, our model assumes that all codons in the genome evolve under purifying selection at the amino acid level: as a result, all amino acid substitutions are considered to be deleterious. In other words, each codon position is assigned either an optimal amino acid given by cognate tRNA pairing with the codon currently observed at that genomic position, or a STOP instruction, such that *j* = 1,…,21. According to Eq. (3), even codons that predominantly produce the optimal amino acid will be penalized if there are non-zero pairing rates for translation into suboptimal amino acids. Similarly, a mutation into a codon for which the rates for translation into suboptimal amino acids are enhanced (for example, mutations of a codon which predominantly produces arginine (Arg) into a predominantly non-Arg codon at a position where the optimal amino acid is Arg) is considered deleterious. Since at each codon position evolutionary dynamics depends on the optimal amino acid, we obtain 21 distinct diagonal matrices containing fitness values for each codon, for 20 amino acids and the STOP instruction (i.e., translating stop codons into amino acids is also considered deleterious in our model).

Equation (1) implements the idea that additional tRNA gene copies should increase the available pool of tRNA molecules which can be paired with the codon *c*, reducing translation times and therefore increasing the fitness of the organism (i.e., as 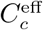 increases, *w*_*j*_(*c*) also increases). However, changes in the tRNA pool may also result in more translation errors, which will be reflected in the increased *s*_*j*_(*c*) (Eq. (3)).

To describe mutations between codons, we have adapted the model of Tamura and Nei (1993). According to this model, *µ*_*c′c*_ = *βπ*_*c′*_, where *µ*_*c′c*_ is the mutation rate per generation from the nucleotide trimer *c* to *c′*, *π*_*c*_ is the steady-state frequency of the nucleotide trimer *c*, and *β* is a scale factor. We compute values of *π_c_* from intergenic sequences which are assumed to evolve under the influence of mutational forces only (Tamura and Nei, 1993; Yang, 2006). The no-selection assumption is supported by the observation that trimeric nucleotide frequencies are very similar in the intergenic regions of all the species we have examined (Fig. S1). Additionally, two transition/transversion rate biases are included when the trimer substitution involves a pyrimidine-to-pyrimidine (*C*↔*T*) exchange (*κ*_1_), or a purine-to-purine (*A*↔*G*) exchange (*κ*_2_). For example, the mutation rate from codon CGT to codon CGC is given by *βκ*_1_*π*_CGC_, whereas the CGA→CGC mutation rate is given by *βπ*_CGC_. Mutation rates corresponding to multiple nucleotide substitutions are set to zero.

Our selection-mutation approach allows us to predict genome-wide codon frequencies through a steady-state population genetics model (Materials and Methods). The major features of the approach are illustrated in Fig. 1 using *Escherichia coli* as an example. Figure 1A shows three initial *E. coli* populations which are genetically identical except for a single codon: one population contains the wild-type codon ATA at position 101 in the thrA gene (position 1 is the start codon), whereas the other two contain codons with single-nucleotide mutations: ATG and AAA, respectively. After a fixed period of time, the three progeny populations have different sizes due to differences in their growth rates (Fig. 1B). The thrA codon under consideration is at a location which, according to the genetic code and the fact that the wild-type codon is ATA, codes optimally for isoleucine (Ile). Figure 1C shows mRNA transcripts produced in the three *E. coli* strains, with colored boxes around codons corresponding to the predominantly translated amino acid in each case: Ile (green), Met (blue), and Lys (red). The lowest-fitness strain has experienced an ATA→AAA mutation, resulting in a codon which cannot be translated into the optimal amino acid, Ile, even through wobble pairing. In comparison, the ATG strain has higher fitness since it can produce Ile through wobble pairing: however, the ATG codon is primarily translated into Met through cognate pairing. The wild-type ATA strain has the highest fitness as it predominantly produces Ile, even though the cognate tRNA of ATA is in fact not present in *E. coli* (Fig. 1D-F).

**FIG. 1.**
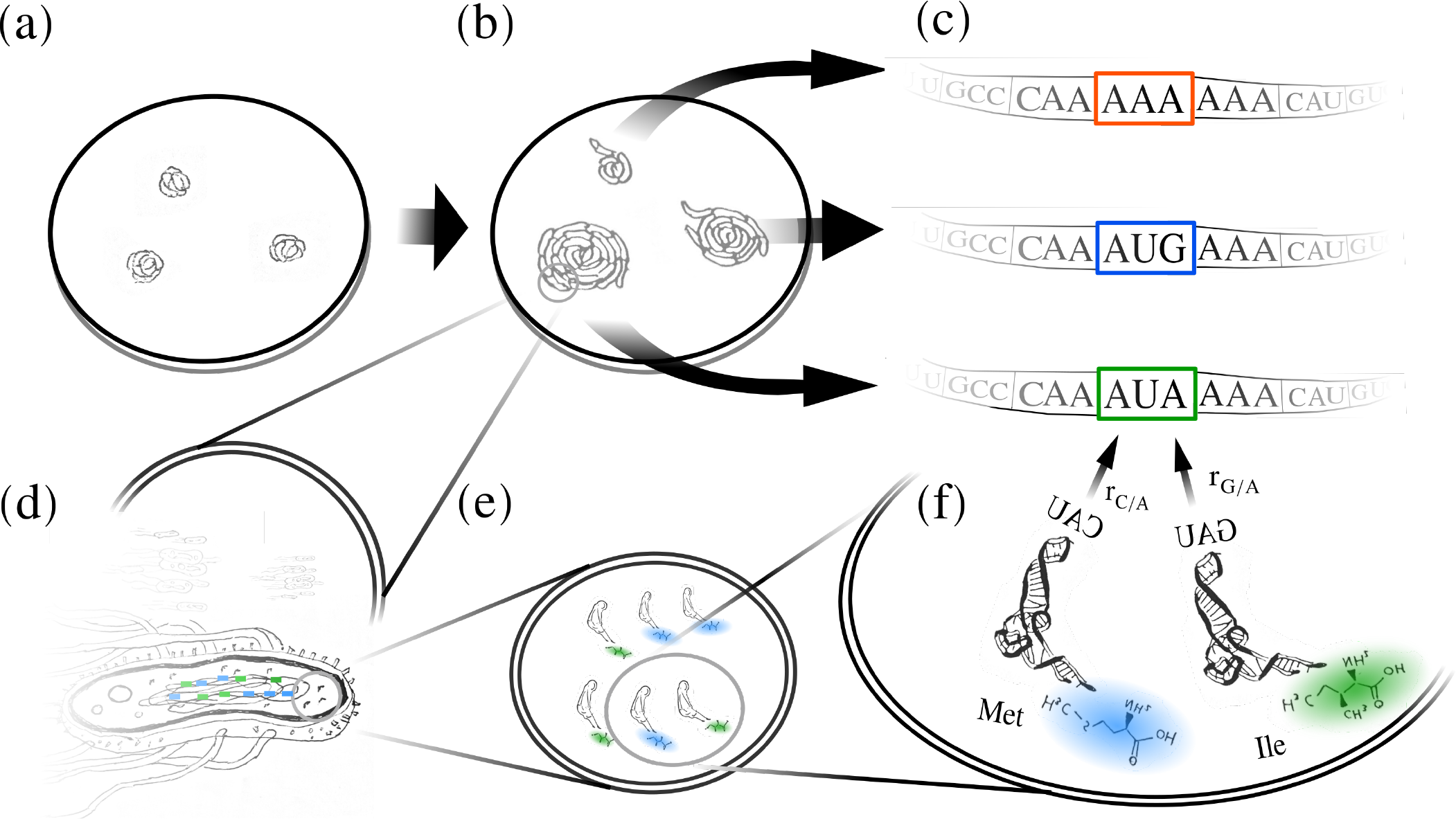
Illustration of the biophysical fitness model on *E. coli* populations. (a) Three initial *E. coli* populations: one wild-type and two with a single-nucleotide mutation (ATA→ATG, ATA→AAA) at codon 101 in the thrA gene. (b) The same *E. coli* populations after a fixed period of growth. (c) RNA transcripts of the thrA gene from all three strains. Each colored box around the codon in question indicates the amino acid that is primarily translated, with green, blue and red corresponding to Ile, Met, and Lys, respectively. (d) An *E. coli* cell with the tRNA gene copies for Ile (green) and Met (blue) shown as colored rectangles. (e) A magnified portion of the cell with three tRNA molecules charged with Ile (green) and three more charged with Met (blue). The proportions of each type of tRNA molecule roughly match the proportions of gene copies in (d), as assumed in our model. (f) A further magnification of (e) with two representative tRNAs shown in molecular detail. The two tRNA molecules shown, one charged with Met and the other with Ile, are present in the K-12 MG1655 *E. coli* and can bind AUA through wobble pairing, with wobble rates *r*_C/A_ and *r*_G/A_, respectively. Note that there is no cognate tRNA for this codon.

### Hierarchy of evolutionary models

To determine which biophysical factors contribute most to the codon bias and what level of detail is necessary to predict genome-wide codon frequencies, we have constructed a hierarchy of models which include from 3 to 19 free parameters (see Table 1 for detailed descriptions), and fit the models to *E. coli* (K-12 MG1655) genomic data. Specifically, each model was fit to minimize the *L*^1^ distance:

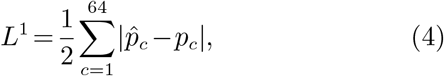

where 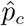 and *p*_*c*_ are predicted and observed genome-wide codon frequencies, respectively (see SI, section S1.1 for a detailed description of the global optimization algorithm). Each model was subjected to 5-fold cross-validation: all genomic codons were randomly divided into 5 subsets of equal size, and the model was fitted separately on each subset, with 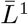 denoting the average *L*^1^ distance resulting from these 5 fits. For the purposes of cross-validation, *L*^1^ distances were computed between codon frequencies predicted by each of the 5 fits and codon frequencies observed in each of the other 4 codon subsets which were not used to fit the model in the current round. The cross-validation score, 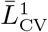, was then computed by averaging first over the other 4 subsets left out of the current fit and, finally, over the 5 independent fits.

**Table 1.**
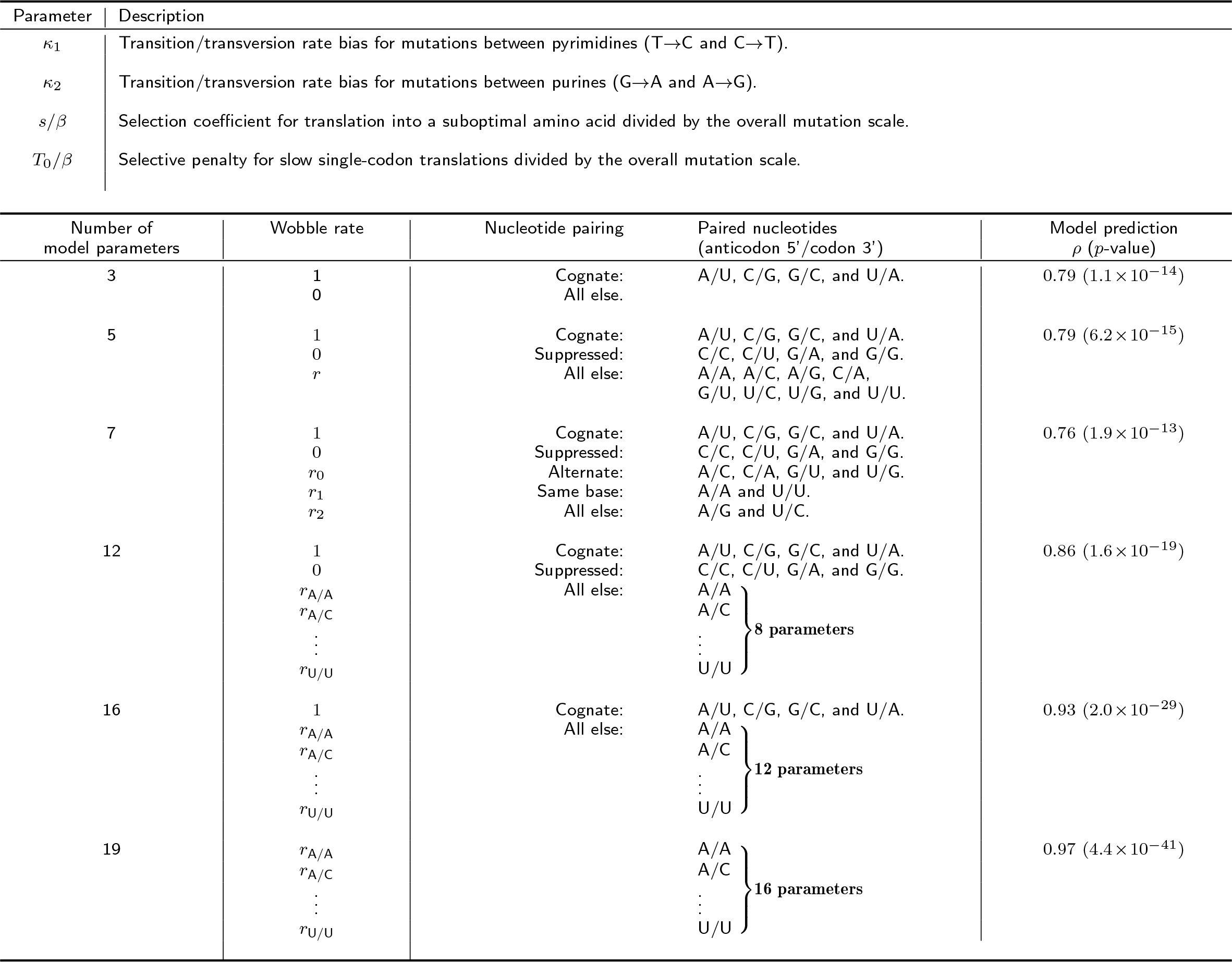
Hierarchy of models fit to *E. coli* genomic data. All models share the same three mutation and selection parameters *κ*_1_, *κ*_2_, and *s/β*, and all except the 3-parameter model also include *T*_0_*/β*. The pairing rates are parameterized according to various categories of nucleotide base pairing: Watson-Crick (Cognate); disallowed according to Murphy IV and Ramakrishnan (2004) (Suppressed); different in type, purine or pyrimidine, and are not disallowed or cognate (Alternate); and two nucleotides of the same type that are not disallowed (Same base). *ρ*: Pearson correlation coefficient between predicted and observed frequencies, *p*: the corresponding p-value.

| Parameter | Description |
| --- | --- |
| $\kappa_1$ | Transition/transversion rate bias for mutations between pyrimidines (T→C and C→T). |
| $\kappa_2$ | Transition/transversion rate bias for mutations between purines (G→A and A→G). |
| $s/\beta$ | Selection coefficient for translation into a suboptimal amino acid divided by the overall mutation scale. |
| $T_0/\beta$ | Selective penalty for slow single-codon translations divided by the overall mutation scale. |

| Number of model parameters | Wobble rate | Nucleotide pairing | Paired nucleotides (anticodon 5'/codon 3') | Model prediction $\rho$ (p-value) |
| --- | --- | --- | --- | --- |
| 3 | 1<br>0 | Cognate:<br>All else: | A/U, C/G, G/C, and U/A. | 0.79 ($1.1 \times 10^{-14}$) |
| 5 | 1<br>0<br>$r$ | Cognate:<br>Suppressed:<br>All else: | A/U, C/G, G/C, and U/A.<br>C/C, C/U, G/A, and G/G.<br>A/A, A/C, A/G, C/A, G/U, U/C, U/G, and U/U. | 0.79 ($6.2 \times 10^{-15}$) |
| 7 | 1<br>0<br>$r_0$<br>$r_1$<br>$r_2$ | Cognate:<br>Suppressed:<br>Alternate:<br>Same base:<br>All else: | A/U, C/G, G/C, and U/A.<br>C/C, C/U, G/A, and G/G.<br>A/C, C/A, G/U, and U/G.<br>A/A and U/U.<br>A/G and U/C. | 0.76 ($1.9 \times 10^{-13}$) |
| 12 | 1<br>0<br>$r_{A/A}$<br>$r_{A/C}$<br>⋮<br>$r_{U/U}$ | Cognate:<br>Suppressed:<br>All else: | A/U, C/G, G/C, and U/A.<br>C/C, C/U, G/A, and G/G.<br>A/A<br>A/C<br>⋮<br>U/U } **8 parameters** | 0.86 ($1.6 \times 10^{-19}$) |
| 16 | 1<br>$r_{A/A}$<br>$r_{A/C}$<br>⋮<br>$r_{U/U}$ | Cognate:<br>All else: | A/U, C/G, G/C, and U/A.<br>A/A<br>A/C<br>⋮<br>U/U } **12 parameters** | 0.93 ($2.0 \times 10^{-29}$) |
| 19 | $r_{A/A}$<br>$r_{A/C}$<br>⋮<br>$r_{U/U}$ | | A/A<br>A/C<br>⋮<br>U/U } **16 parameters** | 0.97 ($4.4 \times 10^{-41}$) |

The first model we have examined is a minimal model which does not consider wobble pairing or the fitness penalty for slow translation and therefore only includes transition/transversion mutational parameters *κ*_1_ and *κ*_2_ and the amino acid selection parameter *s/β*. Note that the codon frequencies are affected only by the ratio of the amino acid selection coefficient *s*, which penalizes translation into suboptimal amino acids, and the overall mutation scale *β* (see SI, section S1.2 for an additional discussion). Under this 3-parameter model, genome-wide codon frequencies are determined by a combination of mutation rate biases and mutational proximity to deleterious sequences (i.e., mutational robustness). We illustrate this point using 6 Arg codons as a representative example (Fig. 2A,B). Under the 3-parameter model there is a marked enrichment of the frequencies of CGC, CGG, and CGT codons and a suppression of AGA and AGG codon frequencies, even though all 6 codons have the same fitness. These trends, with the exception of the CGG enrichment, match genome-wide codon frequency data, and are not present in the intergenic regions (Fig. 2A).

**FIG. 2.**
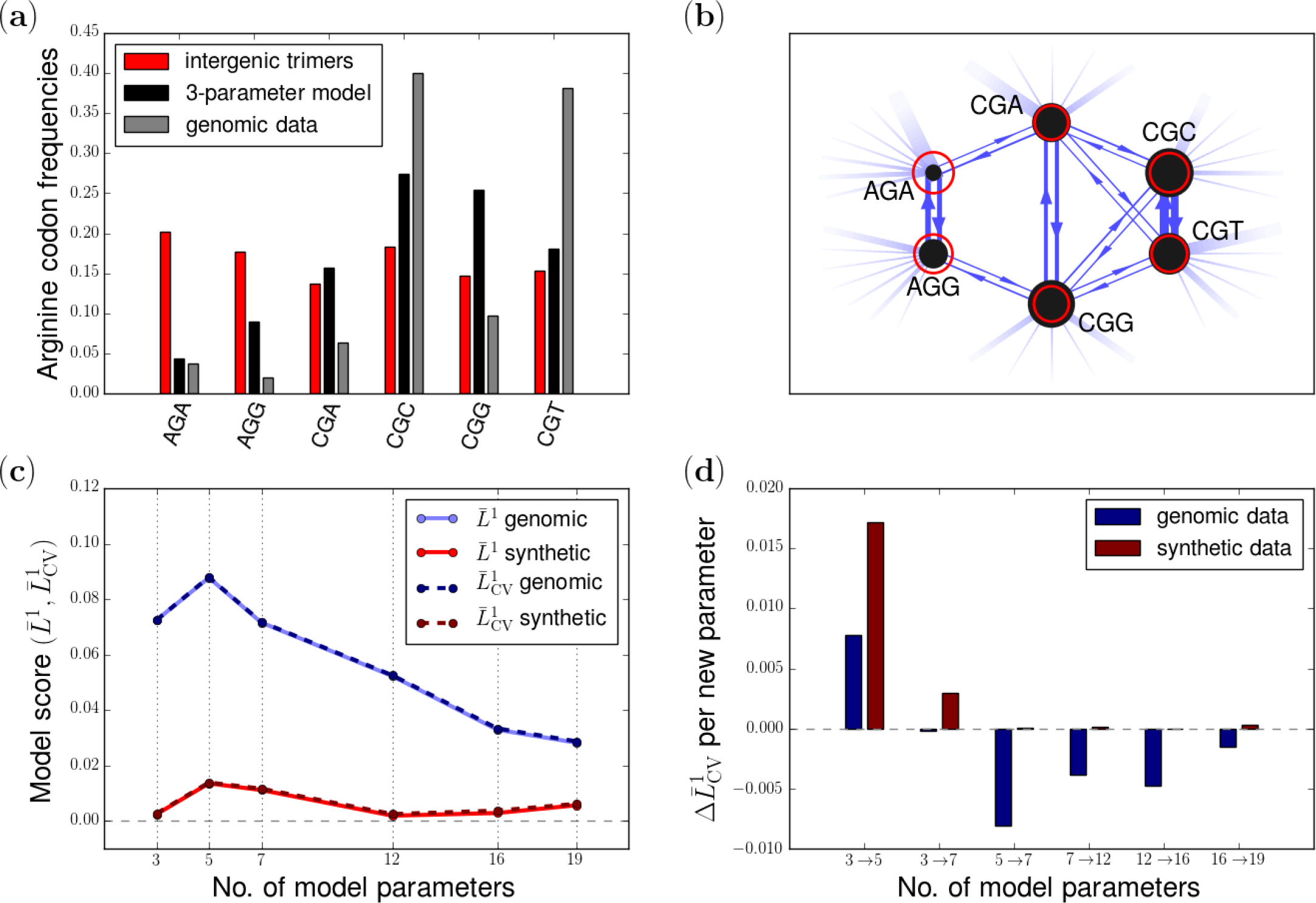
Prediction of genome-wide codon frequencies in *E. coli*. (a) Codon frequencies of the arginine (Arg) group predicted by the 3-parameter model (black) and found in coding regions (grey), and nucleotide trimer frequencies in the intergenic regions (red). (b) The single-point mutational network formed by the codons which translate into Arg according to the standard genetic code. The width of each line is proportional to the mutation rate, with an arrow indicating the direction of mutation. The fading lines represent all mutation rates from Arg to the corresponding non-Arg codons. The size of each circle indicates the frequency at which each codon sequence occurs in intergenic trimers (red) and when mutation and selection against non-Arg codons are taken into account (3-parameter model; black). (c) Model scores 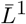 (solid lines) and 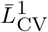(dashed lines) as a function of model complexity. Each model was fit to genomic data (blue lines) and synthetic data (red lines). (d) Normalized difference of 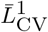 model scores, 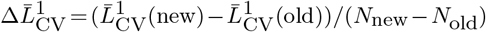, in going from a less complex (“old”) to a more complex (“new”) model. *N* new and *N*_old_ denote the number of model parameters in the old and new models, respectively.

Next, we examined a family of models which in addition to *κ*_1_, *κ*_2_, and *s/β* include a fitness penalty for slow codon translation, *T*_0_*/β*, with *T*_0_ defined in Eq. (1). In addition, each model in the hierarchy includes an increasingly diverse set of pairing rates (Table 1). Specifically, the 5-parameter model has a single parameter describing all non-cognate pairing rates. In this model, cognate pairings are assumed to occur at a rate of *r*_*n′/n*_ = 1, while four pairings are suppressed (*r*_*n′/n*_ = 0) based on the crystallographic analysis of wobble base pairs in the context of the ribosomal decoding center (Murphy IV and Ramakrishnan, 2004). The remaining 8 rates are described by a single free parameter, *r*. The 7- parameter model replaces this single parameter with three rates: *r*_0_, which accounts for pairings across nucleotide types (purine to pyrimidine) expected to be closest to cognate pairing; *r*_1_, which characterizes all same-base pairings that are not already suppressed on the basis of crystallographic evidence; and *r*_2_, which accounts for the two remaining pairings. In the 12-parameter model, rates for 8 pairings that are neither cognate nor suppressed are allowed to vary individually. In the 16-parameter model, the assumption that some of the wobble pairings are suppressed is relaxed, resulting in 4 additional pairing rates. Finally, in the 19-parameter model the assumption that all cognate pairings have a rate of *r*_*n′/n*_ = 1 is relaxed, and each of the possible 16 pairings is assigned an individual rate. Since there is now a degeneracy in the model related to the fact that the 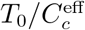 ratio remains invariant in Eq. (1) if both *T*_0_ and all wobble rates are scaled by the same factor, we have chosen to set *T*_0_*/β* = 1, resulting in 19 independent parameters. An alternative approach in which one of the cognate rates was set to 1.0 and *T*_0_*/β* was allowed to vary yielded numerically inferior solutions.

Since 63 independent codon frequencies are fit to models containing from 3 to 19 independent parameters, it is important to ensure that there is no overfitting. Figure 2C demonstrates the quality-of-fit scores 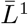 and 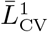 for each of the models described above. A standard way of checking the extent of overfitting, 5-fold cross-validation, has limited applicability here since codon frequencies are very similar in all 5 subsets, as manifested by the high degree of similarity between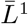 anbd 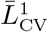 in all Fig. 2C fits. Thus, to investigate the issue of overfitting from a different angle, we have carried out model fits not only on genomic codon frequencies (blue lines), but also on synthetically generated data for which models previously fit on genomic data were used to generate artificial codon counts (for a full description of synthetic data generation, see SI, section S1.1). These counts were then used in a subsequent round of model fitting (red lines). The idea is to provide a score baseline in which a given model type is employed to both generate the synthetic data and carry out subsequent parameter inference. This two-step procedure leads to consistent recovery of all model parameters used in generating the synthetic data (Table S1). As can be seen in Fig. 2C, there is no trend in the model scores of fits on synthetic data as the model complexity increases, and for each model type genomic fit scores are significantly above synthetic fit scores, indicating the absence of overfitting. Furthermore, model scores of fits on genomic data improve with model complexity, suggesting that overall increasing the model complexity is beneficial. Note however that the genomic scores become worse in going from the 3- to 5- parameter model, showing that an increase in the number of model parameters does not necessarily guarantee an improvement in fitting performance.

We have also investigated the effects of intentional overfitting on synthetic data. To this effect, an “old” model with lower complexity, previously fit on genomic data, was used to generate the codon counts, which were subsequently used to fit a “new,” higher-complexity model (red bars in Fig. 2D). Surprisingly, this overfitting always resulted in worse model scores, again underscoring that increasing model complexity does not necessarily lead to better scores, due to both essential differences in model parameterization and the lack of numerical convergence. However, this effect becomes very slight on the higher-complexity end of the model spectrum. To investigate this issue further, we have generated synthetic data using the 7-parameter model, and fit all model types to it (Fig. S2). We observe that, as expected, lower-complexity models are not able to fit the synthetic dataset as well as the “native” 7-parameter model. Furthermore, fitting more complex models does not offer any marked improvements in model scores.

In contrast to the results based on synthetic data, there is a significant improvement in model performance on genomic data with each increase in model complexity (blue bars in Fig. 2D), with a sole exception of the 3- and 5-parameter model pair. However, the gains in model scores diminish gradually, indicating that increasing the number of parameters beyond 19 is unlikely to lead to further significant improvements in model performance. Given that the 19-parameter model yields the best performance, we have chosen it for all further analysis carried out in this study. The model’s predictions in *E. coli* are shown in Fig. 3 and Fig. S3A, indicating that our approach is capable of reproducing all the major features observed in genome-wide codon frequencies in this organism.

**FIG. 3.**
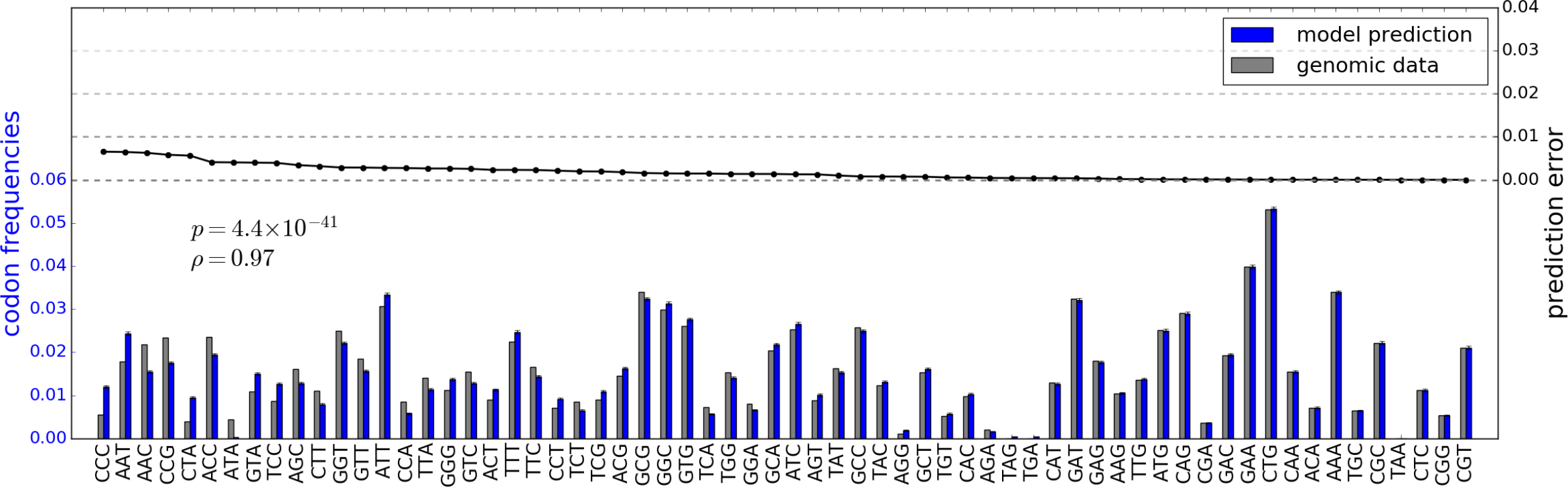
Prediction of codon frequencies in *E. coli*. Codon frequencies predicted using the 19-parameter model (blue), and genome-wide frequencies observed in *E. coli* (grey). All codons are sorted by the absolute magnitude of the prediction error, defined as the absolute magnitude of the difference between predicted, 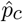, and observed, *p*_*c*_, frequencies of each codon 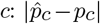. The Pearson correlation coefficient *ρ* between predicted and observed frequencies is also shown, along with the corresponding p-value.

### Modeling codon bias in multiple species

We have fit the 19-parameter model to 20 organisms spanning both unicellular and multicellular life forms (Fig. 4; see SI, section S1.3 for details of genomic data acquisition). In each case, the model fits the data to a high degree of accuracy, with the Pearson correlation coefficients in the [0.80,0.98] range, with an average of 0.91 (Fig. S3B). The inferred biophysical and population genetics parameters and the 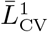 cross-validation scores for each organism are available in Supplementary Dataset S1; distributions of these parameters are summarized in Fig. 5.

**FIG. 4.**
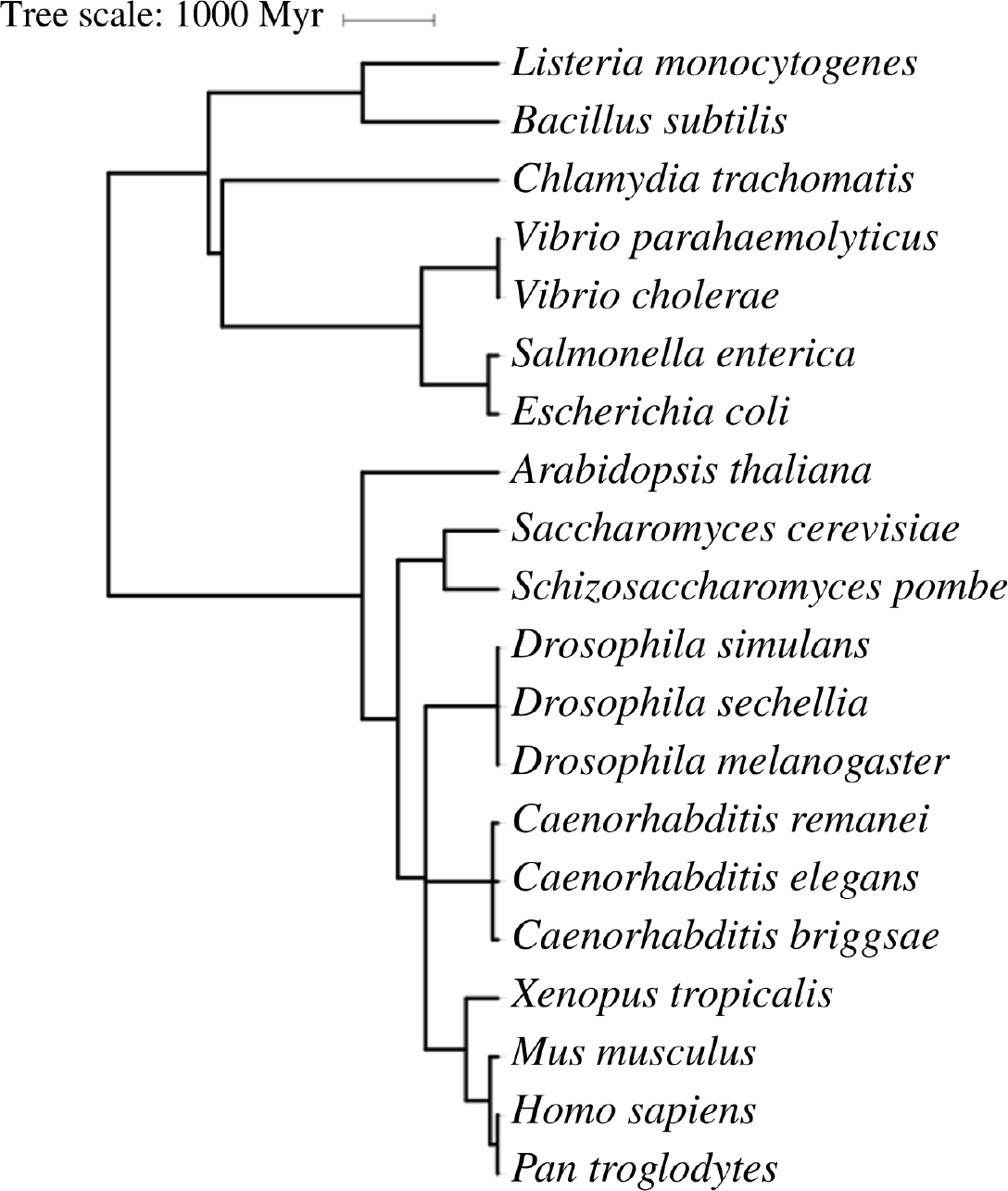
Phylogenetic relationships between all organisms included in this study. The divergence times between all species examined in this study were set to the estimated values reported in the TimeTree database (Hedges *et al.*, 2006; Kumar *et al.*, 2017). These divergence times were then used to construct the phylogenetic tree via the Interactive Tree Of Life (Letunic and Bork, 2016).

**FIG. 5.**
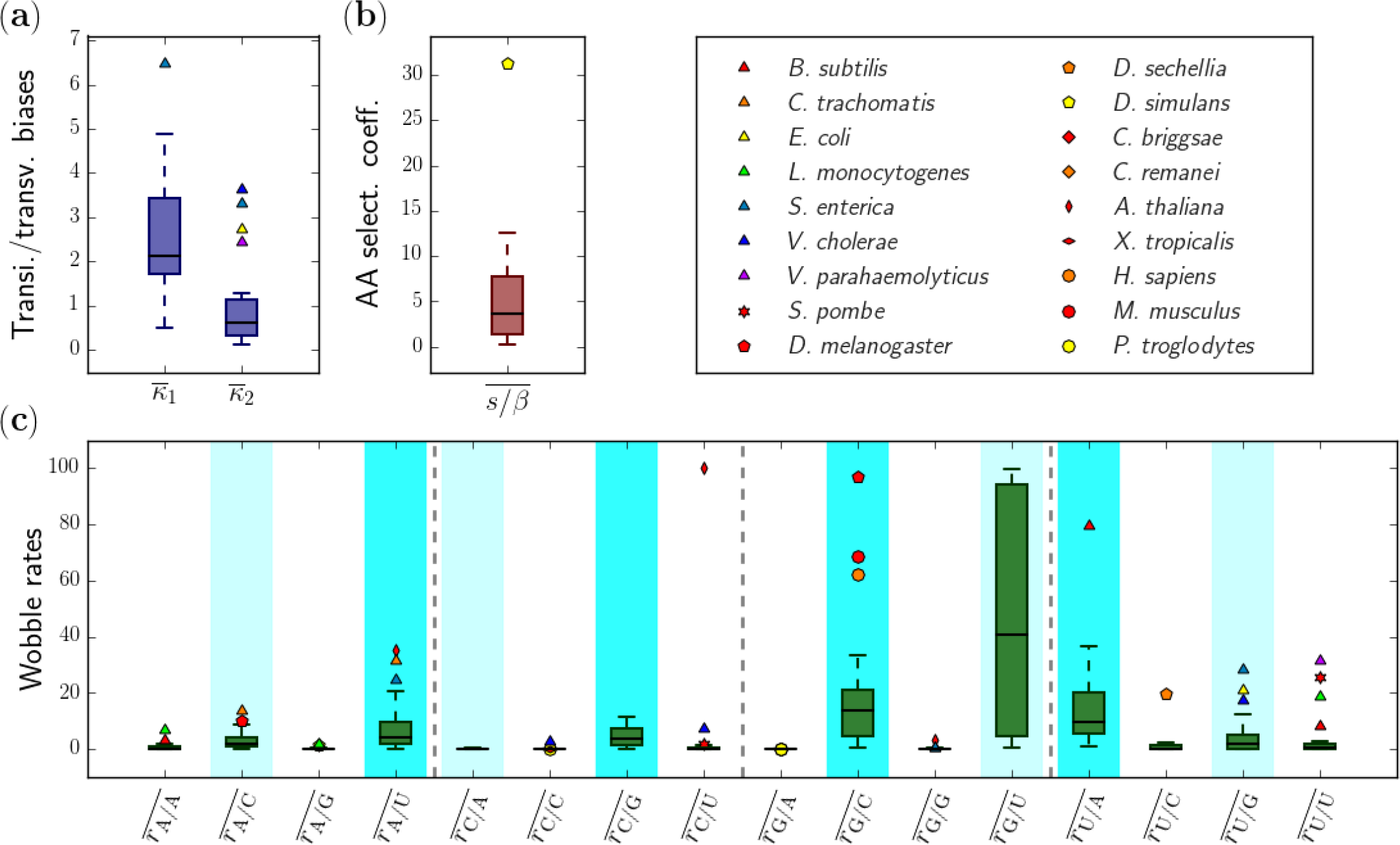
Distributions of inferred biophysical and population genetics parameter values across 20 organisms. All models are fitted separately on 5 codon subsets and the resulting parameters averaged, as indicated by the overbar. For each parameter averaged in this way, median values across all organisms as well as the first, *Q*_1_, and third, *Q*_3_, quartiles are plotted using box-and-whisker plots. The locations of upper and lower whiskers are given by the largest data point below *Q*_3_ +1.5(*Q*_3_*−Q*_1_) and the smallest data point above *Q*_1_− 1.5(*Q*_3_*−Q*_1_). Data points which extend outside of this range are considered outliers and plotted explicitly using species-specific symbols. (a) Transition/transversion rate biases 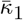 and 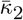, (b) amino acid selection coefficients 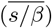, and (c) wobble rates 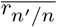. The wobble rates are separated into four sets by vertical dashed grey lines, one for each anticodon nucleotide. The cognate pairings are highlighted in solid cyan, and non-cognate pairings with alternate nucleotide types (purine to pyrimidine pairings) are highlighted in faded cyan.

As might be expected, the values of the two transition/transversion rate biases, *κ*_1_ and *κ*_2_, are fairly conserved, especially in eukaryotes, with larger values generally found in bacteria (Fig. S4, Fig. 5A). This observation is consistent with the fact that trimeric nucleotide frequencies found in intergenic regions, which on average are likely to evolve only under the influence of mutational forces, are nearly organism-independent (Fig. S1). The values of the *κ*_1_ and *κ*_2_ biases are strongly correlated with each other, with *κ*_1_ > *κ*_2_ in all cases. Note that the biases are not always *>* 1, in agreement with a previously reported result (Keller *et al.*, 2007).

We observe strong selection against mis-sense mutations (*s/β* = 5.84 on average), indicating that amino acids translated from the genomic codons on the basis of the standard genetic code are generally optimal and their mutations are deleterious (Fig. 5B). The value of the selection coefficient *s* is closely correlated with the distribution of *s_j_*(*c*) values in each organism (Fig. 6A), indicating that it is a good measure of the strength of selection against amino acid mistranslations. Moreover, the strength of selection for the speed of codon translation is generally weaker than the strength of mutational forces, as measured by the overall mutational scale *β*, although there are also notable exceptions (Fig. 6B). Correspondingly, in the majority of cases selection for mistranslation dominates selection for translation speed (Fig. 6C).

**FIG. 6.**
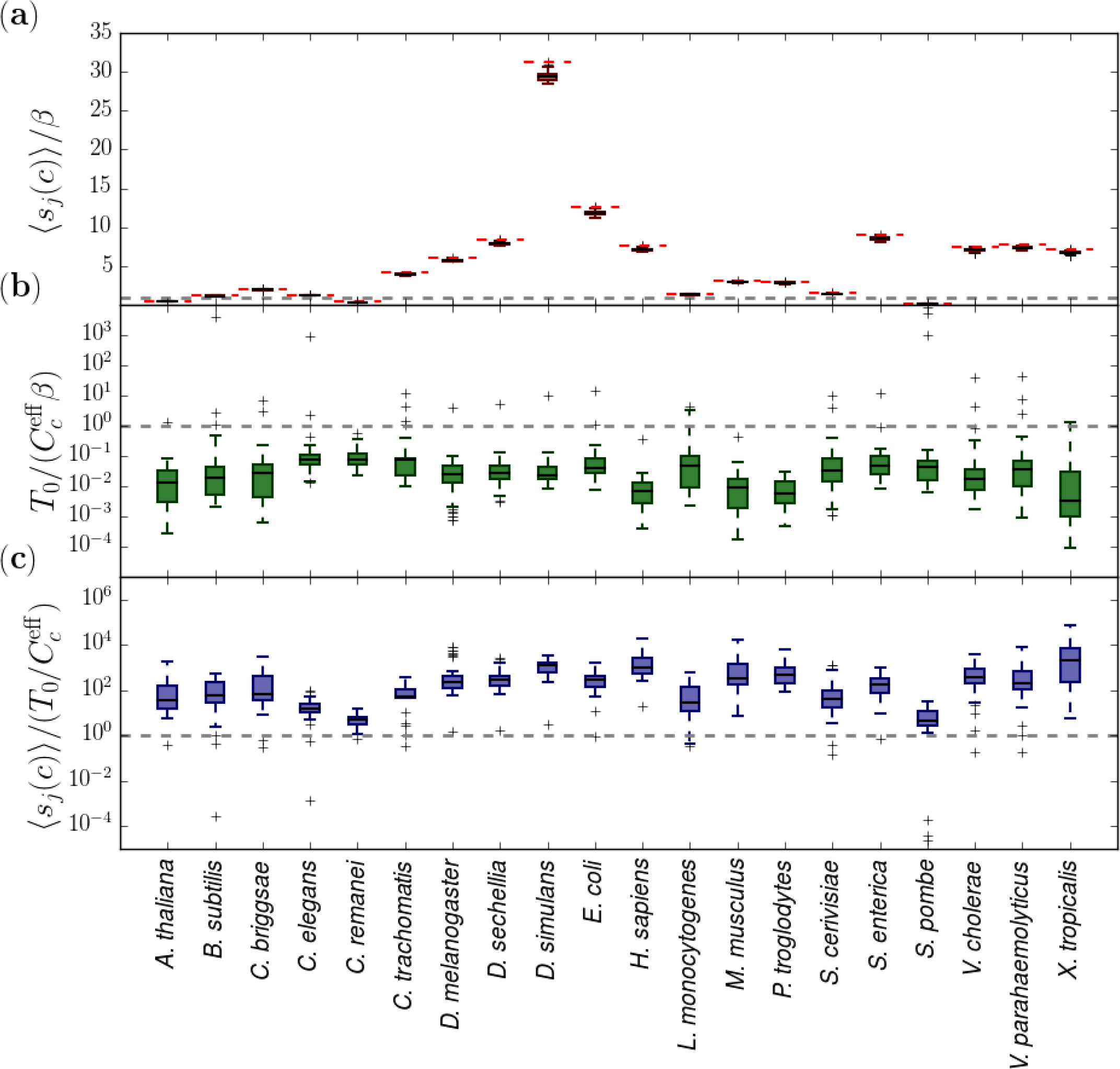
Comparison of selection strengths for speed and accuracy of codon translation. All selection coefficients have been computed by fitting the 19-parameter model to genomic data from 20 organisms. (a) Ratios of the selection coefficients for amino acid mistranslation, *s*_*j*_(*c*) (cf. Eqs. (1) and (3)), averaged over optimal amino acids/STOP instruction as indicated by angle brackets, to the overall mutation scale *β*, shown as box-and-whisker plots for each organism. Horizontal dashed red lines indicate the corresponding value of *s*. (b) Ratios of the selection coefficient for the speed of codon translation, 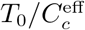 (cf. Eqs. (1) and (2)), to the overall mutation scale *β*, shown as box-and-whisker plots for each organism. (c) Ratios of the two selection coefficients from (a) and (b), shown as box-and-whisker plots for each organism. Horizontal dashed grey lines in panels (a)-(c) indicate where each quantity equals 1.

We have found that in all organisms the rates corresponding to the A/G, C/A, C/C, G/A and G/G pairings are vanishingly small compared to all other rates (Fig. 5C). According to crystallographic evidence (Murphy IV and Ramakrishnan, 2004), C/U, C/C, G/A, and G/G pairings should be sterically disallowed, which is consistent with our findings except for C/U, for which only 3 out of the 20 organisms yield non-vanishing *r*_C/U_ rates: *S. pombe*, *V. Cholerae*, and *A. thaliana*. Additionally, purine-pyrimidine pairings are consistently assigned higher rates than purine-purine and pyrimidine-pyrimidine pairings, with cognate pairings being predominant compared to non-cognate pairings: for example, averages across all species of the 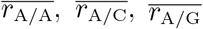, and 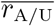 pairing rates are 0.9, 3.3, 0.3, and 8.7, respectively. However, a notable exception is the *r*_G/U_ rate, which is considerably higher than *r*_G/C_ and in fact assumes unrealistically large values for *∼* 50% of the species considered. We do not have a satisfactory explanation for this finding at the moment.

Finally, we have examined a matrix of correlations among 19 model parameters and several additional key values characterizing either the genome (genome size, total number of codons, total number of genes) or the population (effective population size) (Fig. S5). We find that, as expected, the genome size, the total number of codons and the total number of genes are all correlated with each other and anti-correlated with the effective population size and the *κ*_1_, *κ*_2_ mutational biases, the latter observation being consistent with the fact that these biases are higher in prokaryotes (Fig. S4). In contrast, the selection coefficient *s/β* is not strongly correlated with any other parameter, including the effective population size. Finally, we observe that some of the pairing rates (e.g. *r*_A/A_ and *r*_A/G_) are strongly correlated with each other, reducing the effective number of model parameters.

### Estimation of the genome-wide mutation rate

Our biophysical approach has also enabled us to estimate the genome-wide mutation rate per nucleotide per generation as an average over all codon types:

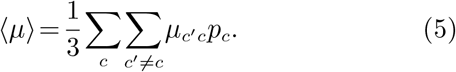

Indeed, following the approach developed by Bulmer (1991), we can estimate *T*_0_ in Eq. (1) directly from the explicit biophysical model of ribosome-mediated translation (see SI, section S1.4 for details). Here, we have focused our attention on *E. coli* and *S. cerevisiae*, for which all the requisite values of biophysical parameters are available in the literature. For both of these organisms, we find that

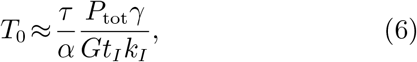

where *P*_tot_ is the total protein production rate in the cell, *G* is the total number of genes, *t*_*I*_ is the average ribosome initiation time, *k*_*I*_ is the average initiation on-rate per free ribosome, and *τ* and *α* are defined in Eqs. (16) and (17), respectively (Materials and Methods). Finally,

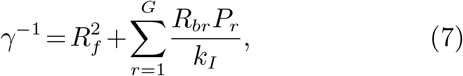

where *P*_*r*_ is the protein production rate of gene *r* (such that 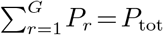), *R*_*br*_ is the average number of ribosomes bound to each transcript of gene *r*, and *R*_*f*_ is the number of free ribosomes in each cell.

Using Eq. (6) and our assumption of *T*_0_*/β* = 1 in the 19- parameter model, we can estimate *β* and, consequently, 〈*µ〉* via Eq. (5), using intergenic trimer frequencies and predicted values of *κ*_1_ and *κ*_2_. Note that although the *T*_0_ = *β* assumption is arbitrary, we estimate *τ/α* from the predicted pairing rates in a way that makes our procedure invariant with respect to rescaling both *T*_0_ and all pairing rates by an arbitrary factor (cf. Eqs. (1), (2), (6), and (S24)). We have estimated the value of *β* using both the most up-to-date data available in the literature and Bulmer’s original data (see Table S2 for input parameter values). Both sets of parameters yield very similar estimates for the average effective mutation rate: 2.4×10^−6^ and 1.1×10^−6^ mutations per nucleotide per generation, respectively. These estimates differ from independent estimates of the genomic mutation rate in *E. coli* (Drake, 1991; Wielgoss *et al.*, 2011), which yield values on the order of 10^−10^ mutations per nucleotide per generation (Table S2). The same calculation in *S. cerevisiae*, which has a similar effective population size (Table S2), has resulted in ⟨*µ*⟩ = 7.1×10^−7^ mutations per nucleotide per generation, which is also higher than the independently estimated mutation rate of 3.3×10^−10^ mutations per nucleotide per generation (Lynch *et al.*, 2008) (see SI, section S1.4 for details and Table S2 for input parameters).

A possible explanation for the observed discrepancy, which is reminiscent of the difficulties encountered by Bulmer in trying to reconcile a population genetics model with the biophysics of mRNA translation (Bulmer, 1991), is that the codon diversity seen in *E. coli* and *S. cerevisiae* genomic data is affected by linkage and may require an explicit treatment of genetic drift, as *µN*_e_ « 1 for both organisms (Table S2). Indeed, genetic drift can contribute to allele diversity observed across multiple sites, even if each individually evolving site is in the monomorphic regime (Manhart *et al.*, 2012; Sella and Hirsh, 2005). Note that our model describes the frequencies of *N*_*c*_ ≃ *G*𝓛/20 = 𝓞(10^5^) codon sites per individual for each fitness landscape, where *L* is the average gene length in codons (494 in *S. cerevisiae* and 319 in *E. coli*), and 20 accounts for the number of distinct amino acid types (positions where the STOP instruction has the highest fitness are excluded from the estimate).

Finally, our analysis yields an inverse relationship between ⟨*µ*⟩ and the total number of genes *G*, which in turn is strongly correlated with the total number of nucleotides in the genome (Fig. S5). This is consistent with Drake’s rule, which states that organisms with larger genomes tend to have smaller mutational rates (Drake, 1991). Multiple-species biophysical data of the type displayed in Table S2 will be required to confirm the trend and estimate its significance quantitatively.

## Discussion

We have developed a population genetics treatment of the biophysical model of codon bias. We assume that genome-wide codon frequencies have reached steady state and model the codon population using a selection-mutation framework in which codons evolve independently of one another. Our model includes a detailed description of codon-level mutations which takes transition/transversion biases into account (Tamura and Nei, 1993; Yang, 2006). Furthermore, there are two kinds of selective forces in the model. We assume that most protein coding regions in the genome evolve under purifying selection and that for each codon, translation into amino acids different from the optimal one (which corresponds to the codon in the standard genetic code) carries a selective penalty. Thus our model incorporates mutational robustness, in which steady-state allele frequencies in a polymorphic population of equal-fitness alleles can be non-uniform, with more robust alleles, separated on average by a higher number of mutational steps from the deleterious alleles, being relatively enriched (van Nimwegen *et al.*, 1999). Interestingly, even the minimal 3-parameter model, which takes only mutation and selection against mistranslation into account and considers only cognate codon-anticodon pairings, is capable of reproducing genome-wide codon frequencies with *ρ* = 0.79 in *E. coli* (Table 1).

In addition to the factors described above, we assume that cellular fitness is proportional to the total protein production rate, which leads to selective penalties for codons with longer translation times. A major factor which determines translation speed is the cellular tRNA concentration, which in our model is assumed to be proportional to the tRNA gene copy numbers in the genome (Kanaya *et al.*, 1999). Finally, codon-anticodon pairing rates are computed on the basis of the wobble hypothesis, such that a mutation in the 3’ nucleotide of a given codon may bring about a complicated set of changes in which the effective tRNA gene copy number may increase or decrease simultaneously with the change in the codon’s mistranslation rate. Thus the final contribution of the codon to the total cellular fitness depends on the delicate balance between speed and accuracy of the codon’s translation, and the genome-wide codon frequencies depend on the steady-state balance between selection and mutation forces. While we have neglected other possible mechanisms of selection on codon usage, such as mRNA toxicity (Mittal *et al.*, 2018), mRNA transcription (Zhou *et al.*, 2016), translation initiation (Bhattacharyya *et al.*, 2018), and co-translational folding (Jacobs and Shakhnovich, 2017), the ability of our model to empirically explain observed patterns of codon usage across many organisms suggests that these mechanisms, while undoubtedly important in some cases, do not play a dominant role in shaping codon usage genome-wide.

We have fit our biophysical model to genomic codon frequencies from 20 organisms. Overall, the model reproduces observed genome-wide patterns of codon usage to a high degree of accuracy (Fig. S3). When codons are ranked based on the accuracy of the model prediction, the codon CTA appears in 8 of the 20 organisms as one of the top 4 least accurately predicted codon frequencies. No such pattern emerges for amino acids. In terms of the predicted model parameters, the values of mutational biases *κ*_1_ and *κ*_2_ are fairly conserved as expected, with larger values typically found in prokaryotes and with *κ*_1_ > *κ*_2_ in all organisms. The universality of mutational rate biases across organisms is consistent with the fact that nucleotide trimer frequencies are strongly conserved in the intergenic regions (Fig. S1).

Furthermore, we observe that codons are under strong selection against mistranslation, with *s/β* = 5.84 when averaged over all organisms (Fig. 5B), and *s/β <* 1 only in *S. pombe*, *C. remanei*, and *A. thaliana*. We have found that in each organism the fitted value of the selective penalty *s*, introduced in Eq. (3), is nearly equal to the mean of the corresponding distribution of the *s*_*j*_(*c*) selection coefficients, defined in Eq. (1) (Fig. 6A). On the other hand, in both *E. coli* and *S. cerevisaie β* is several-fold larger than ⟨*µ*⟩, the genome-wide mutation rate per nucleotide per generation averaged over all codon types (see SI, section S1.4 for details). Thus we expect *s*/⟨*µ*⟩ to be *>* 1 in all organisms, making selection against mistranslation a dominant evolutionary force in comparison with mutational effects.

In contrast, the ratio of the selection coefficient associated with the translation speed to the mutation scale, 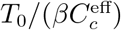, is < 1 on average (Fig. 6B). Thus our model predicts that fitness costs associated with slow translation are often subordinate to the mutational effects, and are much less pronounced than selection against mistranslation (Fig. 6C). Nonetheless, we expect 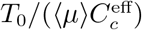 to be ≳ 1 for a nonzero fraction of all codons, indicating that at least in some cases selection against slow translation is an important factor which shapes observed codon frequencies.

Finally, despite the fact that pairing rates are unrestricted in the 19-parameter model, the rates follow well-established patterns consistent with both empirical rules of the wobble hypothesis (Crick, 1966) and atom-level details of codon-anticodon binding on the ribosomal template (Murphy IV and Ramakrishnan, 2004). For example, rates of cognate pairing are much higher than rates of wobble pairing (Fig. 5C), with the sole exception of the *G/U* pairing whose rates are predicted to be anomalously large. Note that in our framework the first two codon positions are assumed to have no effect on the pairing rates.

As an additional test of our approach, we have estimated *T*_0_, defined in Eq. (1), directly using an explicit biophysical model of ribosome-mediated translation originally developed by Bulmer (1991). Bulmer’s model relies on biophysical parameters such as single-codon translation times and translation initiation rates, whose values are available in the literature for *E. coli* and *S. cerevisaie* (Table S2). Estimating *T*_0_ has enabled us to find the average mutation rate per nucleotide, ⟨*µ*⟩, in the coding regions, and compare it with previously published estimates of genome-wide mutation rates (Charlesworth, 2009; Drake, 1991; Lynch *et al.*, 2008, 2011; Wielgoss *et al.*, 2011). Our estimates of ⟨*µ*⟩ are several orders of magnitude higher than the values of *μ* available in the literature. A model of codon evolution which includes genetic drift and linkage between multiple codon loci is necessary to investigate these discrepancies further. Additional refinements of the model could also replace *s* with several fitness penalties which would depend on the physico-chemical similarity of the mistranslated amino acid to the optimal one.

Finally, we note that according to our biophysical framework, *µ* is inversely proportional to the number of genes (Eq. (6)). This is reminiscent of the observation, due to Drake, that organisms with larger genomes tend to have smaller mutational rates (Drake, 1991). We intend to extend our mutation-selection model to all conserved and non-conserved regions of the genome in order to study this correlation in more detail.

## Materials and Methods

### Population genetics model

In order to predict genome-wide codon frequencies, we have employed a mutation-selection population genetics model. We represent codon counts in a population of *N* organisms as a vector with 64 entries, *|N* (*t*)*〉*, and evolve the state of the population from one generation to the next using the deterministic equation:

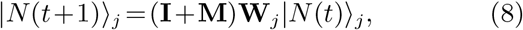

where **W**_*j*_ is a diagonal matrix of fitness values conditioned either on the optimal amino acid or the STOP instruction (i.e., *j* = 1,…,21), **M** is the mutation matrix, and **I** is the identity matrix. The off-diagonal entries of the mutation matrix, **M**_*c′c*_, are the mutation rates from codon *c* to *c′*, and diagonal entries are fixed through Σ_*c*_**M**_*c′c*_ = 0. Equation (8) can be rewritten in terms of the codon frequencies in a population evolving under the same fitness matrix, |*p*(*t*)⟩*_j_* = |*N*(*t*)⟩*_j_*/1|*N* (*t*)⟩*_j_*(|1⟩ is a vector with 1 in every entry),

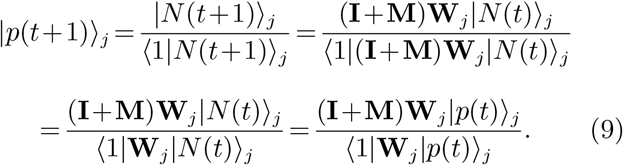

Eventually these frequencies will reach a steady-state |*p*^*ss*^⟩_*j*_ determined by

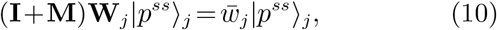

where 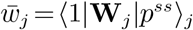 is the average fitness of the corresponding population.

Finally, if each fitness matrix **W**_*j*_ operates at *C*_*j*_ codon locations in the genome, steady-state codon frequencies are given by the genome-wide average:

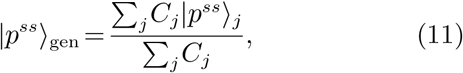

where each |*p*_*ss*_⟩_*j*_ is found using Eq. (10) with the corresponding **W**_*j*_. Note that the mutation rates are assumed to be independent of the fitness matrix *j*, yielding a universal **M** for each species.

### Biophysical model of codon evolution

We model the cell’s fitness, *w*, as proportional to the product of its total protein production rate, *P*_tot_(*c,q,ℓ*), which depends on the presence of codon *c* at location *ℓ* on gene *q* (explicit dependence on all the other codons is suppressed for brevity), and a mistranslation penalty:

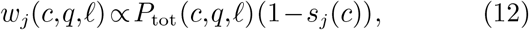

where *s*_*j*_(*c*) is the selection coefficient for codon mistranslation, which we assume to be dependent on the codon’s genomic location only through the optimal amino acid or STOP instruction, *j*, at that location (Eq. (3)).

The change in *P*_tot_ upon mutating the current codon, *c*, at genomic coordinates (*q,ℓ*) into codon *c′* is expected to be small compared to the total protein production rate. The new protein production rate, *P*_tot_(*c′,q,ℓ*), can then be approximated by a first-order expansion,

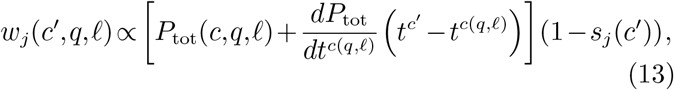

where the single-codon translation time *t*^*c′*^ is assumed to be independent of the codon’s location, and *t*^*c*(*q,ℓ*)^ is the translation time of codon *c* at genomic coordinates (*q,ℓ*).

Next, Eq. (13) is averaged over all codon positions for which *s*_*j*_(*c′*) is the same (that is, over all positions which have the same optimal amino acid or STOP instruction *j* and therefore evolve under the same fitness matrix):

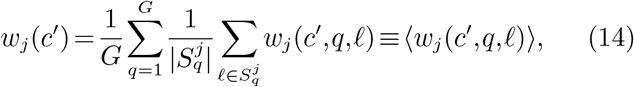

where *G* is the total number of genes, 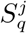 is the set of codon locations with the same optimal amino acid or STOP instruction *j* on gene *q*, and 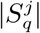 is the number of such locations. Note that all instances for which 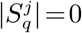 are excluded from the average. We obtain

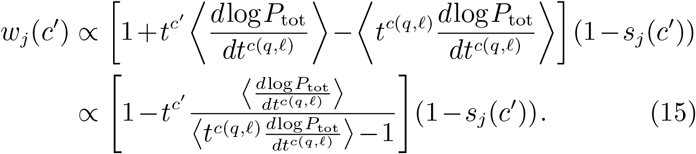

We model the translation time, *t*^*c′*^, as inversely proportional to the tRNA cellular counts:

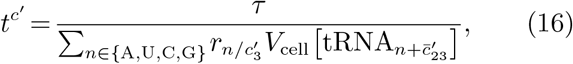

where *V*_*cell*_ is the cell volume, *τ* is the characteristic time scale for tRNA molecules to be acquired by the ribosome for translation, 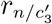 are dimensionless pairing rates at which tRNAs with n as their 5’ anticodon nucleotide bind to the 3’ nucleotide of codon *c′*, denoted 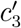 (the other two anticodon nucleotides are always cognate to *c′*), and 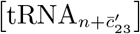 are concentrations of tRNAs with anticodon 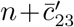 where 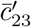 denotes the second and third nucleotides of the reverse complement of *c′*.We assume that the tRNA gene copy number, denoted as 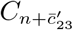, is proportional to the tRNA cellular counts:

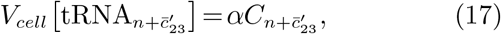

where *α* is a proportionality constant, leading to

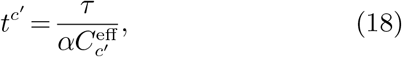

with the effective gene copy number 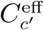 given by Eq. (2).

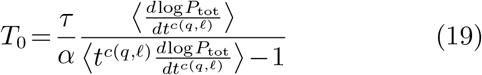

Eq. (15) reduces to Eq. (1). The 64 fitness values for each codon, computed using Eq. (1) and conditioned on the optimal amino acid or STOP instruction *j*, provide the diagonal entries of the fitness matrix **W**_*j*_.

## Acknowledgments

MM was supported by an NIH F32 fellowship (GM116217). AVM and WBK acknowledge financial and logistical support from the Center for Quantitative Biology at Rutgers.

