## Supplementary Material for "Inferring biophysical models of evolution from genome-wide patterns of codon usage"

### S1 Supplementary Materials and Methods

#### S1.1 Numerical determination of model parameters

**Global optimization algorithm.** To fit codon frequencies, we have developed a heuristic optimization algorithm whose objective is to minimize the $L^1$ distance between model predictions and data. The algorithm proceeds as follows:

1. Initialize a set of random model parameters: $\vec{x} = (\kappa_1, \kappa_2, s/\beta, ...)$, where all parameters are drawn from a uniform distribution in the $[0, 10^2]$ range.

2. Generate a list of all possible moves through parameter space, $\vec{x}'_i = \vec{x} + \Delta\vec{x}_i$, in which two changes to parameter values are applied simultaneously as shown below:

$$\begin{aligned}
\vec{x}'_1 &= (\kappa_1 + 2\Delta, \kappa_2, s/\beta, ...), \\
\vec{x}'_2 &= (\kappa_1 - 2\Delta, \kappa_2, s/\beta, ...), \\
\vec{x}'_3 &= (\kappa_1 + \Delta, \kappa_2 + \Delta, s/\beta, ...), \\
\vec{x}'_4 &= (\kappa_1 - \Delta, \kappa_2 + \Delta, s/\beta, ...), \\
&\cdots
\end{aligned}$$

   In each move, wobble rates are enforced to be in the $[0, 10^2]$ range and all other parameters in the $[0, 10^6]$ range, with all moves resulting in out-of-range values excluded from subsequent evaluation. Initially, $\Delta = 10$.

3. From top to bottom of the list, sequentially evaluate the $L^1$ score of each move until $L^1(\vec{x}'_i) < L^1(\vec{x})$ is obtained for $i$th move; accept this beneficial move.

4. If a beneficial move is found in step 3, attempt to move in the direction defined by $\Delta\vec{x}_i = \vec{x}'_i - \vec{x}$ repeatedly until no further gain is observed. Specifically, consider

$$\vec{x}'_{\text{new}} = \vec{x}'_i + m\Delta\vec{x}_i,$$

   where $m$ is initialized to $10^6$ and subsequently reduced by a factor of 10 for each $L^1$ evaluation which does not result in a score improvement. If an improvement is found, the move is accepted and the evaluations continue from the new position, starting with the same value of $m$ that led to the score improvement. The entire process is terminated when $m = 1$ and subsequent evaluations of $\vec{x}'_{\text{new}}$ yield no further gains. The algorithm then goes back to step 3 and continues the search from the next entry in the move set listed in step 2, but with the new $\vec{x}$ resulting from all the beneficial moves accepted during the extended search in the $\Delta\vec{x}_i$ direction (note that $m = 1$ at this point).

5. If no score improvement is found after a full iteration through the set of moves listed in step 2, the step size $\Delta$ is reduced by a factor of 2 and the search procedure described in steps 3 and 4 is repeated. When this reduction results in $\Delta < 10^{-4}$, the step size is reinitialized to the initial value, $\Delta = 10$, and the algorithm restarts from step 2.

6. The run is terminated after $10^3$ evaluations of the $L^1$ score.

Throughout the run, the average rate at which $L^1$ decreases per function evaluation is computed using a list of 25 most recent $L^1$ function evaluations (computations of the average rate commence when at least 25 function evaluations have been performed). This average rate is used to estimate the final expected $L^1$ score by linear extrapolation, given the remaining budget of function evaluations. Note that the minimum $L^1$ score is 0. Thus, if the extrapolated estimate of $L^1$ becomes greater than 0 as a result of accepting the latest (slightly) beneficial move, the algorithm is no longer expected to find an optimal set of parameters in the remaining allotted time. Then the algorithm executes step 5, clears the $L^1$ history list, and proceeds to the next move in the list shown in step 2. This allows the algorithm to avoid repeating moves which result in very small score improvements.

The algorithm described above was independently run $10^3$ times starting from randomized initial conditions, and the set of parameters with the lowest $L^1$ was selected. These parameter values were then uniformly randomized by ±5% to generate another $10^3$ starting points, and the algorithm was run again for $10^3$ evaluations of the $L^1$ score. This process was repeated until no further improvement in the $L^1$ score was observed, at which point the best parameter set recorded throughout the run was reported. A set of representative global optimization runs for our hierarchy of models, using *E. coli* genome-wide codon frequencies as input, is shown in Fig. S6.

**Algorithm validation.** To validate our global optimization procedure, we have generated 5 synthetic data sets with sizes equal to those of the *E. coli* genomic data sets by multinomial sampling of the codon frequencies predicted using model parameters obtained by fitting to *E. coli* genomic data. These synthetically generated counts were then used as input data in a subsequent optimization run. The convergence of the algorithm on synthetic data is demonstrated in Fig. S7 for our hierarchy of models, and the parameters recovered during these subsequent optimization runs are compared with the original parameters in Table S1.

#### S1.2 Exact solution in the two-codon case

In order to gain insight into our biophysical model and its dependence on the various model parameters, here we provide an exact solution for a simplified system which consists of two codons, each of which corresponds to a distinct optimal amino acid. With 2 fitness matrices, Eq. 10 yields

$$\begin{aligned}(\mathbf{I}+\mathbf{M})\mathbf{W}_i|p^{ss}\rangle_i &= \begin{pmatrix}1-\mu_{21} & \mu_{12}\\ \mu_{21} & 1-\mu_{12}\end{pmatrix}\begin{pmatrix}w_1^i & 0\\ 0 & w_2^i\end{pmatrix}\begin{pmatrix}p_1^i\\ p_2^i\end{pmatrix}\\ &= \begin{pmatrix}(1-\mu_{21})w_1^i & \mu_{12}w_2^i\\ \mu_{21}w_1^i & (1-\mu_{12})w_2^i\end{pmatrix}\begin{pmatrix}p_1^i\\ p_2^i\end{pmatrix} = \bar{w}^i\begin{pmatrix}p_1^i\\ p_2^i\end{pmatrix}\end{aligned} \tag{S1}$$

where $w_c^i$ and $p_c^i$ denote the fitness and the steady-state frequency of codon $c \in \{1,2\}$ evolving under fitness matrix $\mathbf{W}_i$ ($i \in \{1,2\}$), respectively, and $\bar{w}^i = w_1^i p_1^i + w_2^i p_2^i$ is the corresponding mean fitness. The steady-state frequencies are then given by

$$p_1^i = \frac{1}{2} - \frac{\mu_{21}w_1^i + \mu_{12}w_2^i}{2\Delta w^i} + \frac{1}{2\Delta w^i}\sqrt{(\Delta w^i)^2 - 2(\mu_{21}w_1^i - \mu_{12}w_2^i)\Delta w^i + (\mu_{21}w_1^i + \mu_{12}w_2^i)^2} \tag{S2}$$

and

$$p_2^i = \frac{1}{2} + \frac{\mu_{21}w_1^i + \mu_{12}w_2^i}{2\Delta w^i} - \frac{1}{2\Delta w^i}\sqrt{(\Delta w^i)^2 - 2(\mu_{21}w_1^i - \mu_{12}w_2^i)\Delta w^i + (\mu_{21}w_1^i + \mu_{12}w_2^i)^2}, \tag{S3}$$

where $\Delta w^i = w_1^i - w_2^i$.

If there are $C_i$ genomic locations evolving under fitness matrix $\mathbf{W}_i$, the genome-wide frequencies for each codon $c$ are given by Eq. 11:

$$p_{c,\text{gen}} = \frac{C_1 p_c^1 + C_2 p_c^2}{C_1 + C_2}. \tag{S4}$$

To examine the dependence of steady-state frequencies on the model parameters $\beta$, $\kappa$, $s$, and $T_0$, the following fitnesses and mutation rates are assumed:

$$\begin{aligned}\mu_{21} &= \beta\kappa\pi_2, \quad \mu_{12} = \beta\kappa\pi_1,\\ w_1^1 &= \left(1 - \frac{T_0}{C_1^{\text{eff}}}\right), \quad w_2^1 = \left(1 - \frac{T_0}{C_2^{\text{eff}}}\right)(1-s),\\ w_1^2 &= \left(1 - \frac{T_0}{C_1^{\text{eff}}}\right)(1-s), \quad w_2^2 = \left(1 - \frac{T_0}{C_2^{\text{eff}}}\right),\end{aligned} \tag{S5}$$

where $\pi_c$ is the steady-state frequency of codon $c$ in the absence of selection. Note that mutations between the two codons are assumed to be transitions. With these specifications, $p_1^1$ in Eq. (S2) becomes

$$\begin{aligned}p_1^1 &= \frac{1}{2} - \frac{\beta\kappa\left(1 - \frac{T_0\pi_2}{C_1^{\text{eff}}} - \frac{T_0\pi_1}{C_2^{\text{eff}}}\right)}{2\left[T_0\frac{C_1^{\text{eff}}-C_2^{\text{eff}}}{C_1^{\text{eff}}C_2^{\text{eff}}} + s\left(1 - \frac{T_0}{C_2^{\text{eff}}}\right)\right]}\\ &+ \frac{1}{2}\sqrt{1 - 2\frac{\beta\kappa\pi_2\left(1-\frac{T_0}{C_1^{\text{eff}}}\right) - \beta\kappa\pi_1\left(1-\frac{T_0}{C_2^{\text{eff}}}\right)(1-s)}{T_0\frac{C_1^{\text{eff}}-C_2^{\text{eff}}}{C_1^{\text{eff}}C_2^{\text{eff}}} + s\left(1-\frac{T_0}{C_2^{\text{eff}}}\right)} + \left[\frac{\beta\kappa\left(1-\frac{T_0\pi_2}{C_1^{\text{eff}}} - \frac{T_0\pi_1}{C_2^{\text{eff}}}\right)}{T_0\frac{C_1^{\text{eff}}-C_2^{\text{eff}}}{C_1^{\text{eff}}C_2^{\text{eff}}} + s\left(1-\frac{T_0}{C_2^{\text{eff}}}\right)}\right]^2}.\end{aligned} \tag{S6}$$

To $\mathcal{O}(\beta)$, $\mathcal{O}(s)$, and $\mathcal{O}(T_0)$, Eq. (S6) simplifies to

$$p_1^1 \approx \frac{1}{2} - \frac{\kappa/2}{\frac{T_0}{\beta}\frac{C_1^{\text{eff}}-C_2^{\text{eff}}}{C_1^{\text{eff}}C_2^{\text{eff}}} + \frac{s}{\beta}} + \frac{1}{2}\sqrt{1 - \frac{2\kappa(\pi_2-\pi_1)}{\frac{T_0}{\beta}\frac{C_1^{\text{eff}}-C_2^{\text{eff}}}{C_1^{\text{eff}}C_2^{\text{eff}}} + \frac{s}{\beta}} + \frac{\kappa^2}{\left[\frac{T_0}{\beta}\frac{C_1^{\text{eff}}-C_2^{\text{eff}}}{C_1^{\text{eff}}C_2^{\text{eff}}} + \frac{s}{\beta}\right]^2}}. \tag{S7}$$

A similar expression can be obtained for $p_1^2$, from which $p_2^1 = 1 - p_1^1$ and $p_2^2 = 1 - p_1^2$ follow by normalization. Note that under this approximation all steady-state frequencies, including the genome-wide frequencies $p_{c,\text{gen}}$ in Eq. (S4), only depend on the ratios $s/\beta$ and $T_0/\beta$.

#### S1.3 Genome Sequences and Annotation

All genomic codon, amino acid, and intergenic nucleotide trimer frequency information was extracted for each species from sequence and annotation data in the GenBank file format, downloaded from the NCBI database on 07.13.2018. tRNA gene copy numbers were obtained from the GtRNAdb database [2, 3]. To compute intergenic trimer frequencies, we have removed all nucleotide sequences corresponding to known features, leaving only DNA segments with no currently known functions.

#### S1.4 Biophysical approach to mutation rate estimation

In order to estimate the average mutational rate $\langle\mu\rangle$ (Eq. (5)), we first need to determine the mutational scale parameter $\beta$. Recall that in the 19-parameter model, we have set $T_0/\beta = 1$ without loss of generality because all wobble rates are free parameters not constrained by normalization. Therefore, using Eq. (19) to estimate $T_0$ automatically determines $\beta$. Specifically, to compute the right-hand side of Eq. (19) we have followed a biophysical approach originally developed by Bulmer [1]. In this approach, ribosome translation kinetics are modeled explicitly under the assumption of steady-state translation of each mRNA transcript. We start by evaluating

$$\left\langle \frac{d\log P_{tot}}{dt^{c(q,\ell)}} \right\rangle = \left\langle \sum_{i=1}^{G} \frac{P_i}{P_{tot}} \frac{d\log P_i}{dt^{c(q,\ell)}} \right\rangle, \tag{S8}$$

where $P_i$ is the protein production rate of gene $i$, $P_{tot} = \sum_{i=1}^{G} P_i$ is the total protein production, $G$ is the total number of genes, and $t^{c(q,\ell)}$ is the translation time of codon $c$ found in gene $q$ at position $\ell$. As defined in Eq. (14), the $\langle \dots \rangle$ average is over all codon positions which evolve under the same fitness matrix. For a single mRNA transcript, the total ribosomal on-rate will be equal to the total ribosomal off-rate, or the rate at which ribosomes complete translation, in steady-state. If the average time for translation initiation for the $i$th gene is $t_{Ii}$, the rate at which translation is completed and proteins are produced is given by $1/t_{Ii}$ per mRNA. If there are $m_i$ mRNA transcripts per cell for gene $i$, the protein production rate for this gene is $P_i = m_i/t_{Ii}$, yielding

$$\frac{d\log P_i}{dt^{c(q,\ell)}} = -\frac{1}{t_{Ii}} \frac{dt_{Ii}}{dt^{c(q,\ell)}}. \tag{S9}$$

Further noting that the initiation time, $t_{Ii}$, is the sum of the time for a single ribosome to bind to the transcript and the time for this ribosome to translate far enough for another ribosome to bind, we obtain

$$t_{Ii} = (k_{Ii} R_f)^{-1} + \sum_{j=1}^{L} t^{c(i,j)}, \tag{S10}$$

where $R_f$ is the number of free ribosomes in the cell, $k_{Ii}$ is the on-rate per free ribosome for gene $i$, and $L$ is the ribosomal footprint. Substitution of Eq. (S10) into Eq. (S9) yields

$$\frac{d\log P_i}{dt^{c(q,\ell)}} = \frac{1}{t_{Ii} k_{Ii} R_f^2} \frac{dR_f}{dt^{c(q,\ell)}} - \frac{1}{t_{Ii}} \delta_{iq} \delta_{\ell \leq L}. \tag{S11}$$

The number of free ribosomes, $R_f$, can be expressed as

$$R_f = R_{tot} - \sum_{i=1}^{G} m_i R_{bi}, \tag{S12}$$

where $R_{tot}$ is the total number of ribosomes in the cell (assumed to be constant), and $R_{bi}$ is the number of ribosomes bound to a single mRNA transcript of gene $r$. If $t_{Ti}$ is the average time for a single ribosome to translate,

$$t_{Ti} = \sum_{j=1}^{\mathcal{L}_i} t^{c(i,j)}, \tag{S13}$$

where $\mathcal{L}_i$ is the total number of codons in gene $i$, then $R_{bi}/t_{Ti}$ is the total ribosomal off-rate, and therefore

$$\frac{R_{bi}}{t_{Ti}} = \frac{1}{t_{Ii}} \;\Rightarrow\; R_{bi} = \frac{t_{Ti}}{t_{Ii}}. \tag{S14}$$

under the steady-state assumption in which the ribosomal on and off rates are equal.

The derivative of $R_f$ in Eq. (S11) is then given by

$$\begin{aligned}\frac{dR_f}{dt^{c(q,\ell)}} &= -\sum_{r=1}^{G} m_r \frac{dR_{br}}{dt^{c(q,\ell)}} = -\sum_{r=1} m_r \left[\frac{1}{t_{Ir}}\frac{dt_{Tr}}{dt^{c(q,\ell)}} - \frac{t_{Tr}}{t_{Ir}^2}\frac{dt_{Ir}}{dt^{c(q,\ell)}}\right] \\ &= -\sum_{r=1}^{G} m_r \left[\frac{\delta_{rq}}{t_{Ir}} - \frac{t_{Tr}}{t_{Ir}^2}\left\{\delta_{rq}\delta_{\ell\le L} - \frac{1}{k_{Ir}R_f^2}\frac{dR_f}{dt^{c(q,\ell)}}\right\}\right] \\ &= (R_{bq}\delta_{\ell\le L} - 1)P_q - \frac{1}{R_f^2}\frac{dR_f}{dt^{c(q,\ell)}}\sum_{r=1}^{G}\frac{R_{br}P_r}{k_{Ir}},\end{aligned} \tag{S15}$$

yielding

$$\frac{dR_f}{dt^{c(q,\ell)}} = \frac{(R_{bq}\delta_{\ell\le L} - 1)P_q}{1 + \frac{1}{R_f^2}\sum_{r=1}^{G}\frac{R_{br}P_r}{k_{Ir}}}, \tag{S16}$$

where $\delta_{\ell\le L} = 1$ if $1 \le \ell \le L$, and 0 otherwise.

Using this result, Eq. (S11) becomes

$$\frac{d\log P_i}{dt^{c(q,\ell)}} = \frac{(R_{bq}\delta_{\ell\le L} - 1)P_q}{\left[R_f^2 + \sum_{r=1}^{G}\frac{R_{br}P_r}{k_{Ir}}\right]t_{Ii}k_{Ii}} - \frac{1}{t_{Ii}}\delta_{iq}\delta_{\ell\le L}. \tag{S17}$$

Finally, Eq. (S8) can be written as

$$\left\langle \frac{d\log P_{tot}}{dt^{c(q,\ell)}}\right\rangle = \left\langle \sum_{i=1}^{G}\frac{P_i}{P_{tot}}\left\{\frac{(R_{bq}\delta_{\ell\le L} - 1)P_q}{\left[R_f^2 + \sum_{r=1}^{G}\frac{R_{br}P_r}{k_{Ir}}\right]t_{Ii}k_{Ii}} - \frac{1}{t_{Ii}}\delta_{iq}\delta_{\ell\le L}\right\}\right\rangle. \tag{S18}$$

To evaluate Eq. (S18) numerically, we have replaced all single-codon translation times and gene-specific initiation rates with typical values, $t^{c(q,\ell)} \to t$ and $k_{Iq} \to k_I$. Equation (S18) now simplifies to

$$\left\langle \frac{d\log P_{\text{tot}}}{dt^{c(q,\ell)}}\right\rangle \approx -\frac{P_{\text{tot}}\gamma}{Gt_Ik_I} + \sum_{q=1}^{G}\frac{P_q(R_{bq}\gamma P_{\text{tot}} - k_I)}{G|S_q^s|t_Ik_IP_{\text{tot}}}\sum_{\ell\in S_q^s}\delta_{\ell\le L}, \tag{S19}$$

where

$$\gamma = \left(R_f^2 + \sum_{r=1}^{G}\frac{R_{br}P_r}{k_I}\right)^{-1}. \tag{S20}$$

**Estimation of the average mutation rate in *E. coli*.** Protein production rates $P_q$ were assumed proportional to the relative cellular abundance of proteins produced from gene $q$ (see Ref. [17] for protein abundance data). Using $t = 0.12$ s, $t_I = 62$ s from the Transimulation Web server ([13]; *E. coli* K-12 MG1655) and $L = 10$ codons ([8]), we obtain $k_IR_f = 1.6 \times 10^{-2}$ initiations per second for each mRNA using Eq. (S10). With $V_{\text{cell}} = 6.7 \times 10^{-16}$ liters per cell ([16]) and a free ribosomal volume density of 500 nM ([11]), the number of free ribosomes is $R_f \simeq 200$. Using our estimate of $k_IR_f$, the initiation on-rate is found to be $k_I = 8.2 \times 10^{-5}$ initiations per ribosome per second for each mRNA.

According to Ref. [13], there are on average 47 proteins produced per mRNA, which has an average lifespan of 7.5 minutes, and 3.6 mRNAs per gene are present in the cell. With $G \simeq 4300$ genes in the *E. coli* K-12 MG1655 reference genome, we obtain

$$P_{\text{tot}} = \frac{\left(3.6\frac{\text{mRNAs}}{\text{gene}}\right) \times \left(47\frac{\text{proteins}}{\text{mRNA}}\right)}{(7.5\text{ mins }) \times \left(60\frac{\text{secs}}{\text{min}}\right)} \times 4300\text{ genes } \simeq 1600\text{ proteins/sec}. \tag{S21}$$

Finally, according to Eq. (S14)

$$R_{br} = \frac{t\mathcal{L}_r}{t_I}, \tag{S22}$$

where $\mathcal{L}_r$ is the number of codons in gene $r$. Then Eq. (S20) yields $\gamma = 1.2 \times 10^{-7}$ and, consequently, the first term on the right-hand side of Eq. (S19) is estimated as $-P_{\text{tot}}\gamma/Gt_Ik_I = -9.0 \times 10^{-6}\ s^{-1}$. Since the value of the second term changes depending on the fitness matrix, this term has been evaluated for all 20 amino acids, with an average value of $\mathcal{O}(10^{-8})\ s^{-1}$ and a standard deviation of $\mathcal{O}(10^{-8})\ s^{-1}$.

Clearly, consistent with the expectation that $T_0$ should be independent of amino acid selection, the first term dominates in *E. coli*, yielding

$$\left\langle \frac{d \log P_{\text{tot}}}{dt^{c(q,\ell)}} \right\rangle \approx -9.0 \times 10^{-6} s^{-1}. \tag{S23}$$

Next, we have focused on estimating the $\tau/\alpha$ prefactor in Eq. (19). We have averaged Eq. (18) over all codons using the tRNA gene copy number data from Refs. [2] and [3], in combination with the pairing rates fit using the 19-parameter model (Table S1), and set this average equal to $t = 0.12$ s ([13]): $t = \sum_c t^c p_c$, resulting in

$$\frac{\tau}{\alpha} = t \left( \sum_c \frac{p_c}{\sum_{n \in \{\text{A,U,C,G}\}} r_{n/c_3} C_{n+\bar{c}_{23}}} \right)^{-1} \simeq 1.2 \text{ s}. \tag{S24}$$

Finally, the denominator in Eq. (19) is approximated by

$$\left\langle t^{c(q,\ell)} \frac{d \log P_{\text{tot}}}{dt^{c(q_\ell)}} \right\rangle - 1 = t \left\langle \frac{d \log P_{\text{tot}}}{dt^{c(q_\ell)}} \right\rangle - 1 \approx -1, \tag{S25}$$

such that the predicted value for the mutation scale is

$$\beta = T_0 \approx \frac{\tau}{\alpha} \frac{P_{\text{tot}} \gamma}{G t_I k_I} = 1.0 \times 10^{-5}. \tag{S26}$$

This result can be used to estimate the average mutation rate per nucleotide per generation:

$$\langle \mu \rangle = \frac{\text{expected \# nucleotide mutations per generation}}{\text{\# nucleotides}} = \frac{\sum_c \sum_{c' \neq c} \mu_{c'c} \mathcal{C}_c}{3 \sum_c \mathcal{C}_c} = \frac{1}{3} \sum_c \sum_{c' \neq c} \mu_{c'c} p_c = 2.4 \times 10^{-6}. \tag{S27}$$

where $\mathcal{C}_c$ is the total number of codons of type $c$ and the mutation rates $\mu_{c'c}$ were computed using $\beta$ as described in the main text. For ease of reference, key biophysical parameters discussed above have been summarized in Table S2.

To examine the robustness of our findings, we have repeated the mutation rate calculation using the values of biophysical quantities originally reported by Bulmer [1] (Table S2). Following the above analysis and using the additional assumption that $P_r = m_r/t_I$ ([1]), where $m_r$ is the average number of mRNAs for gene $r$, we obtain

$$\gamma = \left( R_f^2 + \frac{R_{\text{tot}} - R_f}{k_I t_I} \right)^{-1} \simeq 3.3 \times 10^{-8}, \tag{S28}$$

leading to $-P_{\text{tot}}\gamma/G t_I k_I \simeq 7.3 \times 10^{-6}\ s^{-1}$.

Next, we follow Bulmer in replacing gene-specific average numbers of bound ribosomes with an average over all genes, $R_{br} \to G^{-1} \sum_{r=1}^{G} R_{br}$. This simplification leads to

$$\left\langle \frac{d \log P_{\text{tot}}}{dt^{c(q,\ell)}} \right\rangle \approx -\frac{P_{\text{tot}}\gamma}{G t_I k_I} + \frac{(R_b \gamma P_{\text{tot}} - k_I) L}{G t_I k_I \mathcal{L}}, \tag{S29}$$

where $\mathcal{L} = \frac{1}{G} \sum_r \mathcal{L}_r$ is the average gene length in codons, and the approximation

$$\sum_{q=1}^{G} \frac{P_q}{|S_q^s|} \sum_{\ell \in S_q^s} \delta_{\ell \leq L} \approx P_{\text{tot}} \frac{L}{\mathcal{L}} \tag{S30}$$

has been made. Equation (S29) can be evaluated directly, resulting in

$$\left\langle \frac{d \log P_{\text{tot}}}{dt^{c(q,\ell)}} \right\rangle \approx -8.2 \times 10^{-6}\ s^{-1}, \tag{S31}$$

in close agreement with Eq. (S23). Next, we use the same effective gene copy numbers $C_c^{\text{eff}}$ and codon frequencies $p_c$ as before to find $\tau/\alpha = 0.56\ s$ via Eq. (S24). This leads to $\beta = 4.6 \times 10^{-6}$ and $\langle \mu \rangle = 1.1 \times 10^{-6}$, close to the estimate in Eq. (S27) which employed more up-to-date parameters.

**Estimation of the average mutation rate in *S. cerevisiae*.** For baker's yeast, an estimate of $R_f = 2.8 \times 10^4$ ribosomes per cell was obtained by taking 15% ([18]) of $R_{\text{tot}}$, reported to be $18.7 \times 10^4$ ribosomes in Ref. [15]. Next, Eq. (S10) was used with $t = 0.10$ s ([6]), $L = 10$ codons ([7]), and $t_I = 54$ s ([13]) to find $k_I = 6.7 \times 10^{-7}$ initiations per ribosome per mRNA per second. According to Ref. [15], $P_{\text{tot}} = 1.3 \times 10^4$ proteins per second, and $G \simeq 6000$ genes in the *S. cerevisiae* reference genome downloaded from NCBI. These values were used in conjunction with the *S. cerevisiae* protein abundance data from Ref. [17] to find $\gamma = 6.2 \times 10^{-11}$ (Eq. (S20)). The first term on the right-hand side of Eq. (S19) was estimated to be $-P_{\text{tot}}\gamma/Gt_I k_I = -3.7 \times 10^{-6}\ s^{-1}$, and once again was found to dominate the second term. Finally, just as before, Eq. (18) was used in combination with the wobble rates predicted by the 19-parameter model to yield $\tau/\alpha = 2.6\ s$, resulting in $\beta = 9.7 \times 10^{-6}$ (Eq. (S26)) and $\langle\mu\rangle = 7.1 \times 10^{-7}$ mutations per nucleotide per generation (Eq. (S27)). For ease of reference, key biophysical parameters have been summarized in Table S2.

#### Supplementary Figures

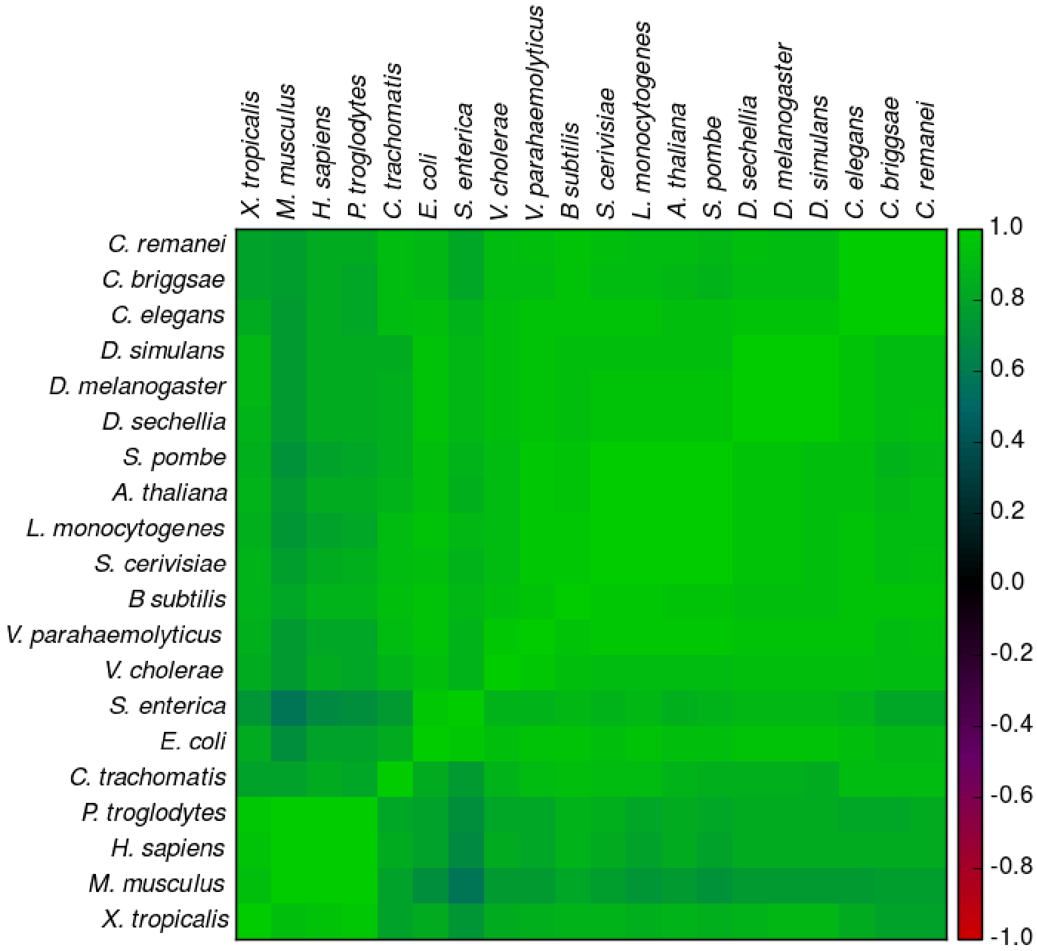


Figure S1: **Covariance matrix of nucleotide trimeric frequencies in intergenic regions for all pairs of organisms considered in this study.** Each entry in this matrix shows the Pearson correlation coefficient between pairs of species-specific trimer frequencies. The lowest entry in this matrix, with the Pearson correlation coefficient $\rho = 0.57$, corresponds to the *S. enterica* – *M. musculus* pair.

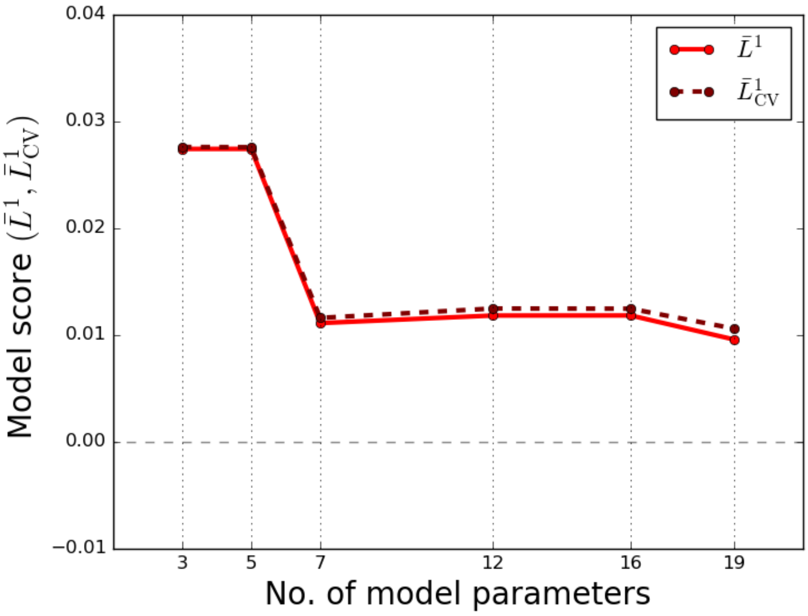


Figure S2: **Model scores for all model types in the hierarchy fitted to synthetic data generated by the 7-parameter model.** Fitting scores $\bar{L}_1$ (solid lines) and cross-validation scores $\bar{L}^1_{\rm CV}$ (dashed lines) are shown as a function of model complexity. All model parameters in the 7-parameter model used to generate the synthetic data were set to values previously found in fitting the model to the codon frequencies from the *E. coli* genome (Table S1).

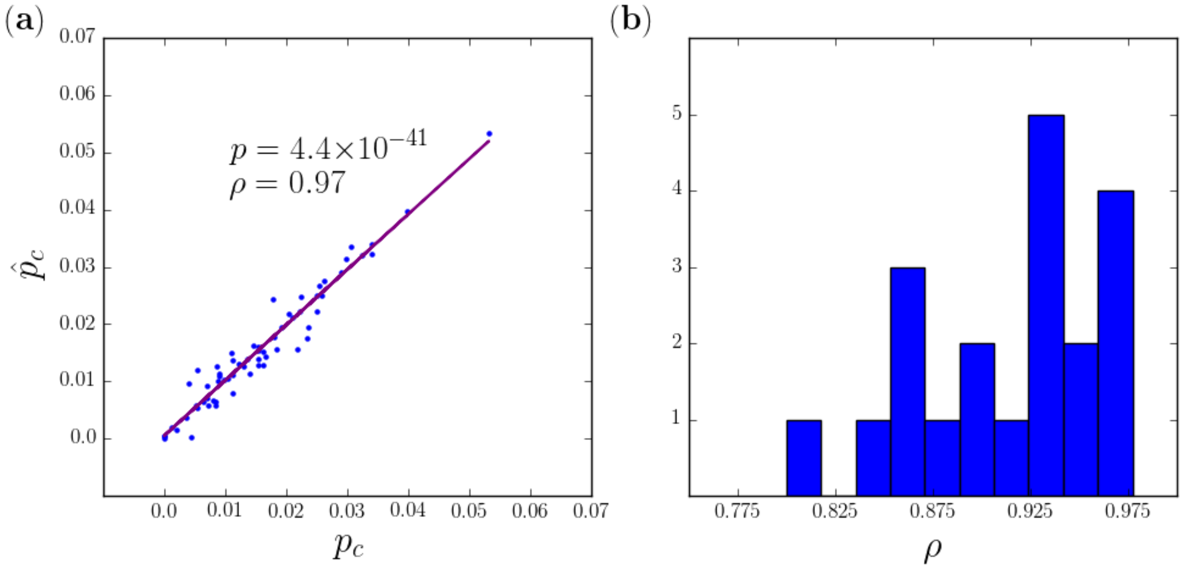


Figure S3: **19-parameter model performance on genomic data.** (a) Predicted versus genomic codon frequencies for *E. coli*, with the Pearson correlation coefficient and the corresponding p-value. (b) Distribution of Pearson correlation coefficients between predicted and observed genome-wide codon frequencies for all 20 species included into this study (Fig. 4).

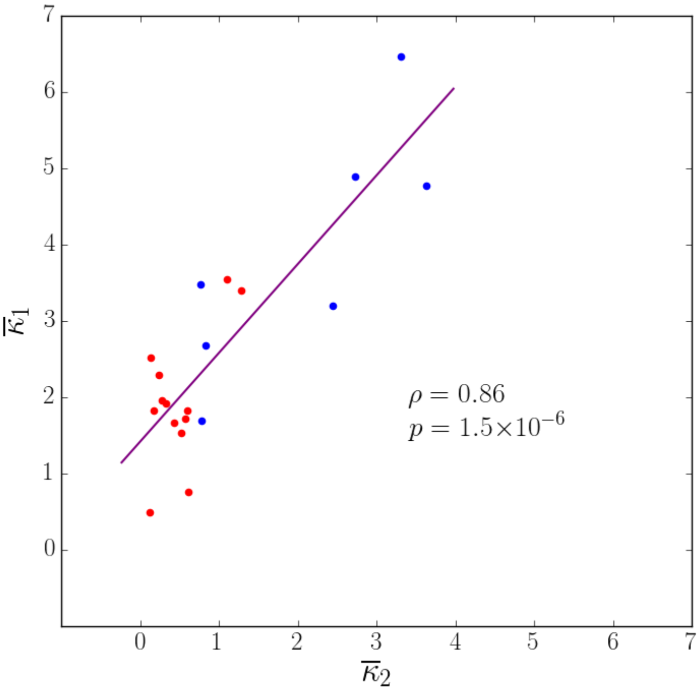


Figure S4: **Correlation between transition/transversion rate biases.** Transition/transversion rate bias parameters $\bar{\kappa}_1$ and $\bar{\kappa}_2$ inferred by fitting the 19-parameter model to genome-wide codon frequencies from 20 species and averaged over 5 distinct subsets of codons. Blue dots: prokaryotes, red dots: eukaryotes. Also shown are the Pearson correlation coefficient $\rho$ and the corresponding p-value.

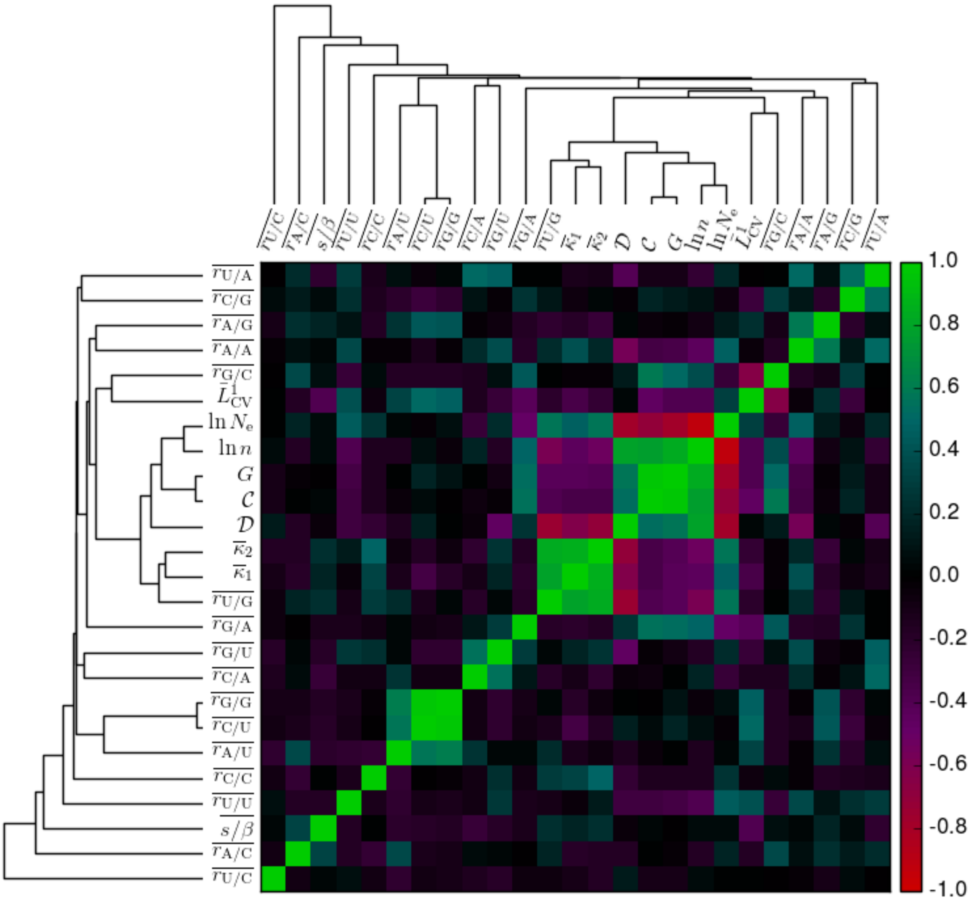


Figure S5: **Covariance matrix between various model and additional parameters**. Each entry in the matrix shows the Pearson correlation coefficient between a pair of parameters, with each parameter available for all 20 species included into this study (Fig. 4). In addition to the 19 model parameters (Table 1), the total number of codons, $\mathcal{C}$, genome size in nucleotides, $n$, effective population size, $N_\mathrm{e}$, the cross-validation model score, $\bar{L}^1_\mathrm{CV}$, the total number of genes, $G$, and the domain label, $\mathcal{D} \in \{\mathrm{Bacteria}, \mathrm{Eukarya}\}$, are included. $N_\mathrm{e}$ estimates have been obtained from Refs. [4] (*C. elegans*, *C. remanei*, *D. melanogaster*, *E. coli*, *H. sapiens*, *P. troglodytes*), [14] (*C. briggsae*), [10] (*B. subtilis*), [12] (*M. musculus*), [5] (*S. enterica*), and [9] (*A. thaliana*, *C. trachomatis*, *D. sechellia*, *D. simulans*, *L. monocytogenes*, *S. cerevisiae*, *S. pombe*, *V. cholerae*, *V. parahaemolyticus*, *X. tropicalis*). All parameters were hierarchically clustered using the linkage package from the SciPy Hierarchical clustering library with the 'single' method and default settings.

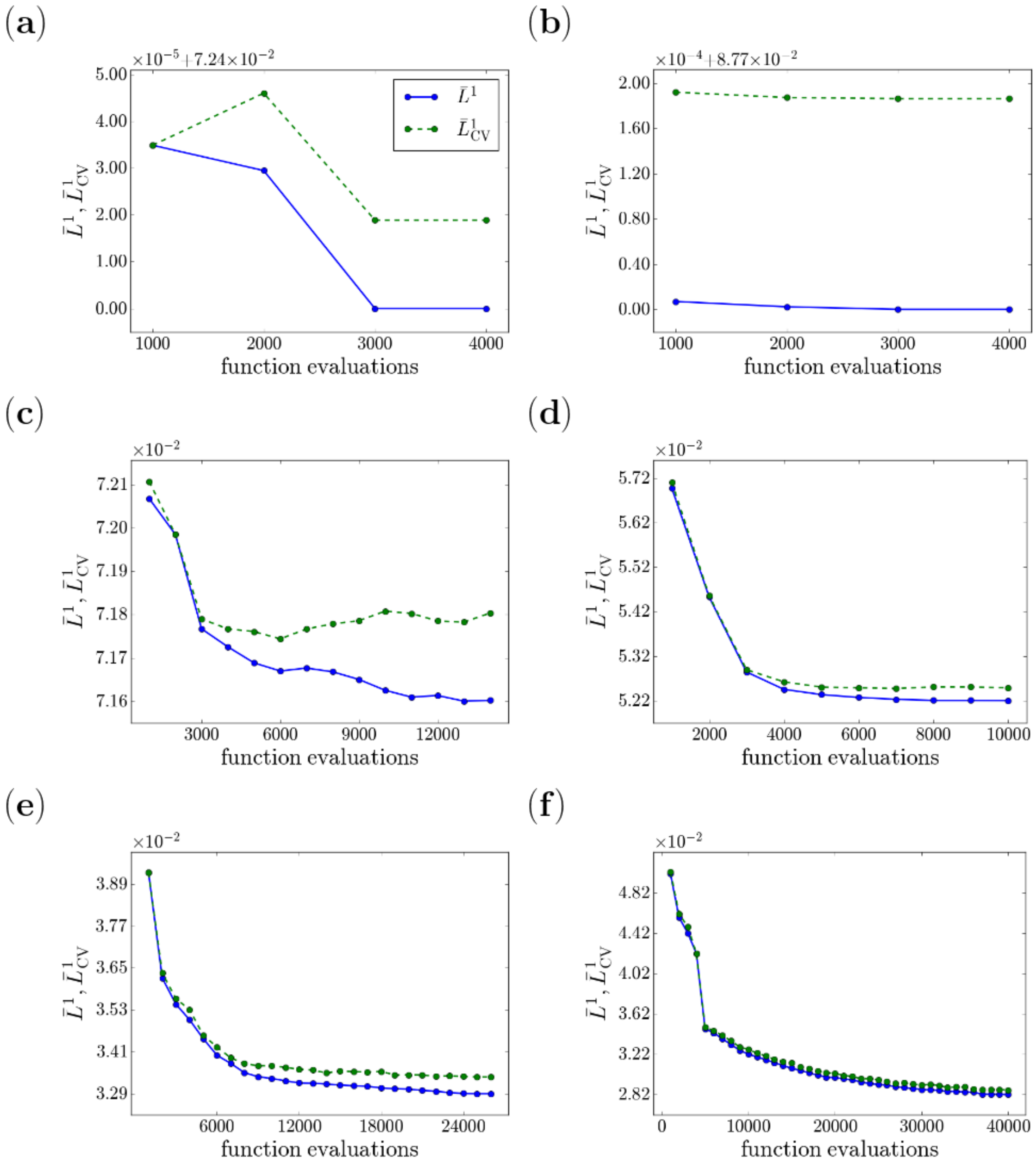


Figure S6: **Convergence of the algorithm fitted to the codon frequencies from the *E. coli* genome.** Model scores $\bar{L}_1$ (solid lines) and $\bar{L}^1_{\mathrm{CV}}$ (dashed lines) are shown vs. the total number of function evaluations in the 3-parameter (a), 5-parameter (b), 7-parameter (c), 12-parameter (d), 16-parameter (e), and 19-parameter (f) models.

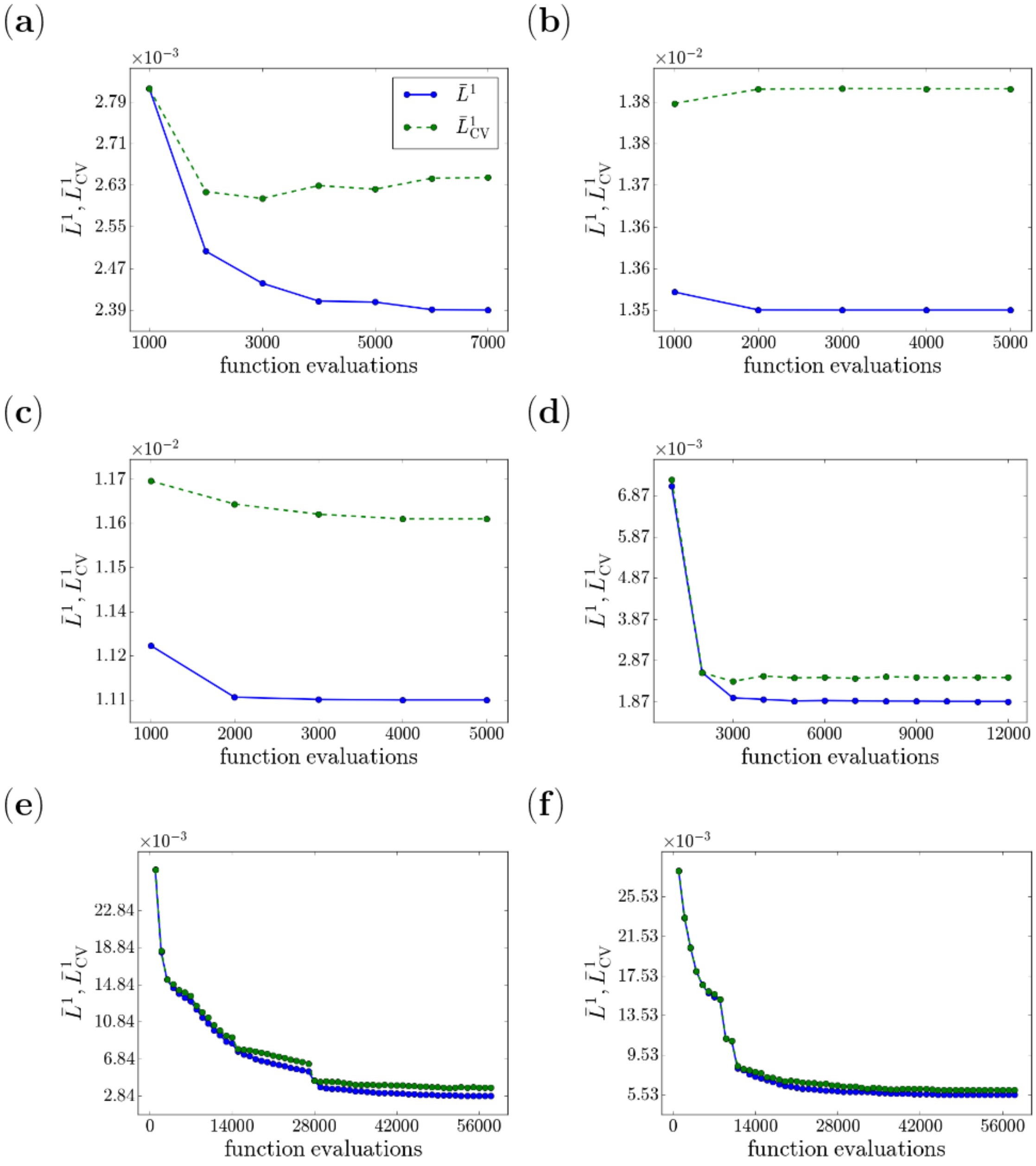


Figure S7: **Convergence of the algorithm fitted to the codon frequencies based on synthetic datasets.** Model scores $\bar{L}_1$ (solid lines) and $\bar{L}^1_{\mathrm{CV}}$ (dashed lines) are shown vs. the total number of function evaluations in the 3-parameter (a), 5-parameter (b), 7-parameter (c), 12-parameter (d), 16-parameter (e), and 19-parameter (f) models. Each fit was performed on synthetic data generated by the same model, with all the model parameters set to values previously found in fitting the model to codon frequencies from the *E. coli* genome (Table S1).

### Supplementary Tables

Table S1: **Sets of parameters used to generate synthetic datasets, and subsequent parameter predictions.** Each parameter set was obtained by fitting the corresponding model to *E. coli* genomic data (first row in each model subsection). Second and third rows show the average and the RMS deviation of model parameters obtained by re-fitting the same model on 5 synthetic datasets which were independently generated using the values in the first row.

3-parameter model

| $\kappa_1$ | $\kappa_2$ | $s/\beta$ |
|---|---|---|
| 4.18 | 3.44 | 644 |
| 4.16 | 3.42 | $2.00 \times 10^5$ |
| .0478 | .0502 | $3.87 \times 10^5$ |

5-parameter model

| $\kappa_1$ | $\kappa_2$ | $s/\beta$ | $T_0/\beta$ | $r$ |
|---|---|---|---|---|
| 3.38 | .211 | .581 | .237 | .659 |
| 3.41 | .214 | .633 | .230 | .650 |
| .0901 | .0192 | .0105 | $2.62 \times 10^{-3}$ | $1.45 \times 10^{-3}$ |

7-parameter model

| $\kappa_1$ | $\kappa_2$ | $s/\beta$ | $T_0/\beta$ | $r_0$ | $r_1$ | $r_2$ |
|---|---|---|---|---|---|---|
| 6.59 | 4.74 | 868 | .0702 | .702 | .000 | .000 |
| 6.65 | 4.76 | $1.39 \times 10^4$ | .0712 | .708 | .000 | .000 |
| .222 | .160 | 7360 | $1.54 \times 10^{-3}$ | .0151 | .000 | .000 |

12-parameter model

| $\kappa_1$ | $\kappa_2$ | $s/\beta$ | $T_0/\beta$ | $r_{A/A}$ | $r_{A/C}$ | $r_{A/G}$ | $r_{C/A}$ |
|---|---|---|---|---|---|---|---|
| 5.23 | 2.85 | 644 | .139 | .216 | .490 | .000 | .000 |
| 5.23 | 2.85 | 8980 | .139 | .209 | .487 | $3.00 \times 10^{-3}$ | .000 |
| .0553 | .0357 | 6480 | $1.25 \times 10^{-3}$ | $7.42 \times 10^{-3}$ | .0124 | $5.56 \times 10^{-3}$ | .000 |

| $r_{G/U}$ | $r_{U/C}$ | $r_{U/G}$ | $r_{U/U}$ |
|---|---|---|---|
| 1.01 | .000 | .605 | .000 |
| 1.00 | .000 | .603 | .000 |
| .0145 | .000 | .0105 | .000 |

16-parameter model

| $\kappa_1$ | $\kappa_2$ | $s/\beta$ | $T_0/\beta$ | $r_{A/A}$ | $r_{A/C}$ | $r_{A/G}$ | $r_{C/A}$ |
|---|---|---|---|---|---|---|---|
| 5.47 | 3.49 | 22.8 | .0997 | .114 | .348 | .000 | $1.22 \times 10^{-5}$ |
| 5.45 | 3.46 | 20.5 | .0999 | .114 | .355 | $1.96 \times 10^{-3}$ | $6.10 \times 10^{-5}$ |
| .0898 | .0950 | 1.39 | $1.40 \times 10^{-3}$ | $4.20 \times 10^{-3}$ | $8.07 \times 10^{-3}$ | $2.85 \times 10^{-3}$ | $1.22 \times 10^{-4}$ |

| $r_{C/C}$ | $r_{C/U}$ | $r_{G/A}$ | $r_{G/G}$ | $r_{G/U}$ | $r_{U/C}$ | $r_{U/G}$ | $r_{U/U}$ |
|---|---|---|---|---|---|---|---|
| .000 | $9.05 \times 10^{-5}$ | $1.10 \times 10^{-3}$ | .0126 | 1.48 | $5.40 \times 10^{-4}$ | 1.26 | .000 |
| .000 | $1.40 \times 10^{-4}$ | $1.18 \times 10^{-3}$ | .0138 | 1.46 | $6.20 \times 10^{-4}$ | 1.24 | .000 |
| .000 | $8.18 \times 10^{-5}$ | $1.94 \times 10^{-4}$ | $1.28 \times 10^{-3}$ | 0.0527 | $4.32 \times 10^{-5}$ | .0385 | .000 |

19-parameter model

| $\kappa_1$ | $\kappa_2$ | $s/\beta$ | $r_{A/A}$ | $r_{A/C}$ | $r_{A/G}$ | $r_{A/U}$ | $r_{C/A}$ |
|---|---|---|---|---|---|---|---|
| 4.90 | 2.73 | 12.6 | 1.08 | 1.68 | $8.06 \times 10^{-3}$ | 2.38 | 8.80 |
| 4.92 | 2.78 | 8.41 | 1.13 | 1.72 | .0127 | 2.42 | 6.81 |
| .0650 | .0272 | .301 | .0751 | .0459 | $8.51 \times 10^{-3}$ | .0878 | $1.81 \times 10^{-3}$ |

| $r_{C/C}$ | $r_{C/G}$ | $r_{C/U}$ | $r_{G/A}$ | $r_{G/C}$ | $r_{G/G}$ | $r_{G/U}$ | $r_{U/A}$ |
|---|---|---|---|---|---|---|---|
| $1.80 \times 10^{-3}$ | 4.26 | .000 | .000 | 18.6 | .313 | 12.0 | 8.19 |
| .000 | 4.37 | $8.17 \times 10^{-5}$ | $1.92 \times 10^{-3}$ | 18.5 | .470 | 12.0 | 8.11 |
| .000 | .157 | $1.63 \times 10^{-4}$ | $1.95 \times 10^{-3}$ | .834 | .0381 | .207 | .122 |

| $r_{U/C}$ | $r_{U/G}$ | $r_{U/U}$ |
|---|---|---|
| .0559 | 21.3 | .000 |
| .0838 | 20.5 | .000 |
| .0117 | 1.03 | .000 |

Table S2: **Summary of key quantities used in mutation rate estimation.** See subsection S1.4 for parameter definitions, references, and additional details.

| | $t$ | $t_I$ | $L$ | $R_f$ | $k_I$ | $P_{\text{tot}}$ | $R_{\text{tot}}$ | $G$ | $N_{\text{e}}$ | $\mu$ |
|---|---|---|---|---|---|---|---|---|---|---|
| This study | | | | | | | | | | |
| *E. coli* | 0.12 | 62 | 10 | 200 | $8.2 \times 10^{-5}$ | 1600 | 42000 | 4300 | $2.5 \times 10^{7}$ | $\sim 10^{-10}$ |
| *S. cer.* | 0.10 | 54 | 10 | $2.8 \times 10^{4}$ | $6.7 \times 10^{-7}$ | $1.3 \times 10^{4}$ | $18.7 \times 10^{4}$ | 6000 | $10^{7}$ | $3.3 \times 10^{-10}$ |
| Bulmer [1] | | | | | | | | | | |
| *E. coli* | 0.056 | 2.0 | 10 | 2800 | $3.6 \times 10^{-4}$ | 680 | 18700 | – | – | – |
